# Dual quorum-sensing control of purine biosynthesis drives pathogenic fitness of *Enterococcus faecalis*

**DOI:** 10.1101/2024.08.13.607696

**Authors:** Soumaya Zlitni, Sierra Bowden, Hila Sberro, Marcelo D. T. Torres, Joan M Vaughan, Antonio F M Pinto, Yishay Pinto, Daniel Fernandez, Hannes Röst, Alan Saghatelian, Cesar de la Fuente-Nunez, Ami S. Bhatt

**Affiliations:** Department of Genetics, Stanford University, Stanford, CA, USA; Department of Medicine (Hematology, Blood and Marrow Transplantation), Stanford University, Stanford, CA, USA; Machine Biology Group, Departments of Psychiatry and Microbiology, Institute for Biomedical Informatics, Institute for Translational Medicine and Therapeutics, Perelman School of Medicine, University of Pennsylvania; Philadelphia, Pennsylvania 19104, USA; Departments of Bioengineering and Chemical and Biomolecular Engineering, School of Engineering and Applied Science, University of Pennsylvania; Philadelphia, Pennsylvania 19104, USA; Department of Chemistry, School of Arts and Sciences, University of Pennsylvania, Philadelphia, Pennsylvania 19104, USA; Penn Institute for Computational Science, University of Pennsylvania; Philadelphia, Pennsylvania 19104, USA; Clayton Foundation Laboratories for Peptide Biology, Salk Institute for Biological Studies, San Diego, CA, USA; Program in Chemistry, Engineering, and Medicine for Human Health (ChEM-H), Stanford University, Stanford, CA 94305, USA; Sarafan ChEM-H Macromolecular Structure Knowledge Center, Stanford University, Stanford, CA 94305, USA; Department of Molecular Genetics, Donnelly Centre for Cellular and Biomolecular Research, The University of Toronto, Toronto, ON, Canada

## Abstract

*Enterococcus faecalis* is a resident of the human gut, though upon translocation to the blood or body tissues, it can be pathogenic. Here we discover and characterize two peptide-based quorum-sensing systems that transcriptionally modulate de novo purine biosynthesis in *E. faecalis*. Using a comparative genomic analysis, we find that most enterococcal species do not encode this system; *E. moraviensis*, *E. haemoperoxidus* and *E. caccae*, three species that are closely related to *E. faecalis*, encode one of the two systems, and only *E. faecalis* encodes both systems. We show that these systems are important for the intracellular survival of *E. faecalis* within macrophages and for the fitness of *E. faecalis* in a murine wound infection model. Taken together, we combine comparative genomics, microbiological, bacterial genetics, transcriptomics, targeted proteomics and animal model experiments to describe a paired quorum sensing mechanism that directly influences central metabolism and impacts the pathogenicity of *E. faecalis*.

## INTRODUCTION

Quorum-sensing (QS) is a communication mechanism employed by many bacterial species to coordinate their behavior based on population density. This allows unicellular microbes to display collective behavior and adapt to environmental challenges more effectively^1,2^. QS is mediated by diffusible chemical signals, ranging from small molecules to small peptides, that are produced by bacteria and accumulate in their environment as they multiply. Once a threshold concentration of the signaling molecule is reached, the bacteria can sense it, and this triggers a coordinated transcriptional response^1,2^. QS has been implicated in various biological functions including virulence^3,4^, biofilm formation^5,6^, motility^7^, sporulation^8,9^, conjugation^10,11^, competence^12^, bioluminescence^13^, and biosynthesis of secondary metabolites, such as bacteriocins^14,15^.

Over the past decades, small molecule-based QS, most common in Gram-negative bacteria, has been studied extensively. By contrast, peptide-based QS, which predominates in Gram-positive bacteria, has garnered less attention. One of the main reasons that peptide-based QS systems are less well understood is that the signaling microproteins, which are encoded by small open reading frames (smORFs), are overlooked in both computational and experimental annotation of microbial genomes^16–19^. Small genes are difficult to distinguish from random in-frame genomic fragments and thus gene prediction tools have classically included a minimum ORF length cutoff. Furthermore, experimental approaches such as Tn-mutagenesis fail to disrupt small genes because of their size, and biochemical identification using targeted and high throughput methods is also challenging. Recent advancements in computational identification and annotation of these prokaryotic smORFs^20–22^, such as the new prokaryotic smORF annotation pipeline smORFinder^23^, have revealed many more microproteins in bacterial genomes than previously known. Among these newly annotated smORFs, a subset likely represent yet undiscovered and uncharacterized peptide-based quorum sensing signals.

There are two main types of peptide-based communication systems in Gram-positive bacteria^24^: systems in which the peptide quorum signal binds to a cell surface receptor, and those in which it binds to an intracellular receptor. In the first type (cell surface receptor system), a small protein precursor is expressed, and as it is secreted, its N-terminal signal sequence is cleaved off, resulting in secretion of the remaining peptide. This mature signaling peptide then interacts with a membrane-bound cell-surface receptor, triggering a cascade of events that ultimately regulates target gene expression. In the second (intracellular receptor system), a small protein precursor is expressed and trafficked extracellularly through its N-terminal signal sequence. The signal sequence is cleaved off, producing a mature peptide that is linear, unmodified, and typically 5-10 amino acids in length^25^. In some cases, the mature QS peptide is further modified by processes such as cyclization. The mature QS peptide is then imported back into the cell through an ATP-binding cassette transporter called oligopeptide permease (Opp). Once inside the cell, it binds to its cytoplasmic receptor, which acts as a transcriptional regulator. The peptide-bound transcriptional regulator can then bind to specific DNA sequences and modulate expression of its target gene(s).

One of the best studied intracellular receptor types of systems is the RRNPP family of signaling systems, named after the peptide receptors of prototypical systems in this family: Rap, Rgg, NprR, PlcR, and PrgX^25,26^. The RRNPP family of communication systems shares characteristic features in their genomic organization, where a typical system consists of a small ORF located directly next to a larger gene that encodes its cognate receptor. The receptors always contain one or multiple C-terminal tetratricopeptide repeats (TPR) or a TPR-like domain. With the exception of the Rap proteins^27^, RRNPP receptors also contain a N-terminal helix-turn-helix type DNA-binding domain^25^. Known RRNPP systems mediate quorum regulation of a broad range of biological processes^25^. For example, Rap systems regulate sporulation and competence in *Bacillus subtilis*^27,28^, PlcR systems regulate virulence in the *Bacillus cereus* group^29^, NprR systems regulate necrotrophism and sporulation in the *Bacillus cereus* group^30,31^, and ComR systems regulate competence in different *Streptococcus spp*^32^. Most known RRNPP systems are present as a single copy within a genome - small peptide and receptor pair - that controls a specific function, as noted above. In *Enterococcus faecalis*, a commensal resident of the human gut and an opportunistic pathogen that is among the leading causes of hospital-acquired infections^33^, there are no known RRNPP systems encoded on the bacterial chromosome. The only two characterized RRNPP systems in this organism are encoded on plasmids, and they control the conjugative transfer of plasmids CF10 and AD1^34^.

In most cases, a single RRNPP system is encoded within a given bacterial genome. *Streptococcus pyogenes* is an exception, as it encodes two very similar systems - the Rgg2/3 systems - within the same organism^35^. Research spanning a series of manuscripts on this system and its function suggests that it is involved in regulating cell surface attributes^25,35–37^. It remains unclear why both systems exist and are widely conserved across all strains of *S. pyogenes*; however, it has been postulated that having two systems that regulate the same downstream function might offer functional redundancy, cooperativity, or provide fine-tuned regulation of downstream processes^38–41^. Taken together, peptide-based QS systems control a range of downstream functions, and there is limited but exciting evidence that duplication and divergence of these systems within an organism can lead to nuanced regulation functions critical for bacterial fitness. Thus, the identification and characterization of such systems, where two related QS programs control the same downstream process, would be particularly exciting.

In this study, we sought to identify novel peptide-based communication systems in bacteria of the human microbiome. To achieve this, we first searched for the minimal elements of a peptide-based communication system: a TPR domain-containing protein and a neighboring smORF. We identified 125 unique systems in 1,661 Human Microbiome Project reference genomes^42^, most of which are of unknown function. Detailed characterization was performed to de-orphan the function of two homologous systems identified in the chromosome of the gut bacterium and opportunistic pathogen, *Enterococcus faecalis.* Furthermore, an evolutionary analysis was conducted to provide insight into when these systems emerged in the evolutionary history of enterococci. Finally, genetic, biochemical, cell biological, and animal studies were carried out to mechanistically dissect the functions of this intriguing set of QS systems.

## RESULTS

### Discovery of new peptide-based quorum sensing systems in human associated microbes

RRNPP family communication systems typically contain a pair of genes: (i) a ‘receptor’ sequence that contains one or more TPR domains and (ii) a microprotein sequence with a signal peptide that, when cleaved, results in a C-terminal short peptide that can bind to and activate the receptor (Figure 1a). We sought to discover new communication systems that contained these genomic features. To search for such putative QS systems in human-associated microbes, we mined all annotated protein-coding genes in 1,661 reference genomes from the Human Microbiome Project (HMP) based on the following criteria: (i) the small open reading frame (smORF) must be ≤ 50 amino acids long, (ii) the smORF must have a N-terminal signal peptide sequence that targets the protein for secretion, (iii) the putative ‘receptor’ gene up- or down-stream of the smORF must encode a protein with at least one TPR domain. Since not all TPR domain-containing proteins interact with microproteins, we began by annotating the putatively secreted smORFs in the HMP genomes to narrow our computational search space before searching for putative TPR domain-containing receptors in the vicinity of the smORFs.

**Figure 1.**
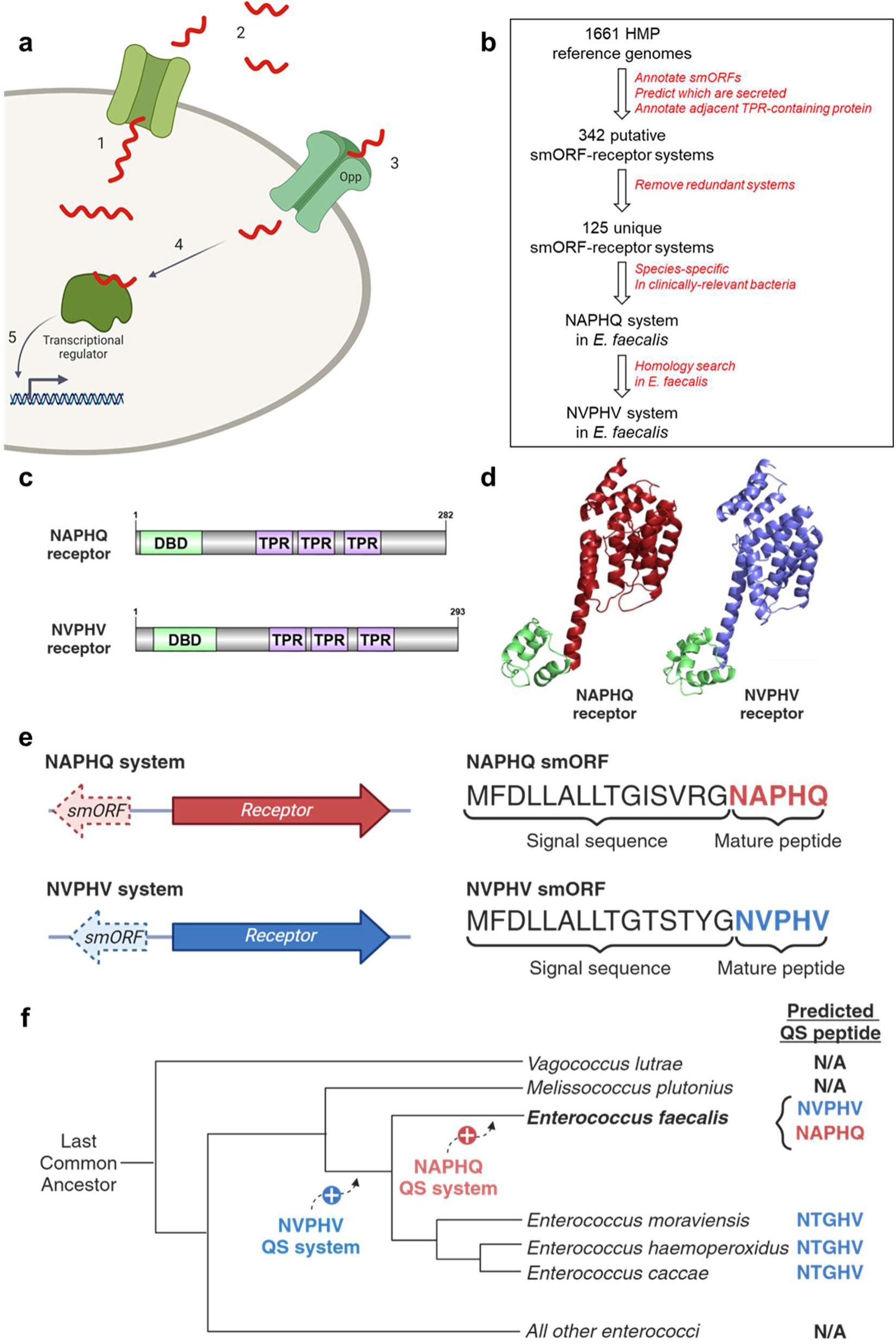
Discovery of novel peptide-based communication systems in HMP reference genomes. **a) RRNPP general mechanism.** Intracellular peptide-based communication systems in Gram-positive bacteria. In this system, (1) a small protein precursor is expressed and secreted extracellularly through its signal sequence. (2) During secretion, it undergoes proteolytic cleavage and sometimes post-translational modification to produce the mature form of the signal peptide. (3) The mature peptide is imported back into the cell through the ATP-binding cassette transporter, oligopeptide permease (Opp). (4) Inside the cell, the signaling peptide binds to its cognate receptor and transcriptional regulator which, in turn, (5) modulates the expression of its target genes. **b) Computational mining for novel peptide communication systems in HMP.** We analyzed a collection of 1661 reference genomes from the Human Microbiome Project (HMP) looking for putative smORF-RRNPP receptor pairs of genes. We identified 125 unique pairs of which two are specific and prevalent in *Enterococcus faecalis* genomes. **c) Putative receptor domain structure of the novel systems.** Similar to most known receptors from the RRNPP family of peptide-based communication systems, the receptors of both systems have a N-terminal DNA-binding domain (DBD; green) and multiple tetratricopeptide (TPR) domains (purple), degenerate 34 amino acid tandem repeats that mediate protein-protein interactions. **d) AlphaFold predicted structures of receptors.** The predicted structures of the NAPHQ receptor (average pLDDT 95.45) and the NVPHV receptor (average pLDDT 96.61). **e) Discovery of two novel peptide-based communication systems in *E. faecalis*.** We identified two putative peptide communication systems in *E. faecalis*, each comprising a large gene encoding a putative receptor/transcription factor and a 20-amino acid small protein. Based on *in silico* prediction of signal sequences, the small proteins are predicted to have a 15-amino acid signal sequence, suggesting that the mature active peptides of these systems are pentapeptides, NAPHQ and NVPHV. **f) Evolutionary analysis of the prevalence of the QS systems in Enterococcal species** Cladogram summarizing the prevalence of RRNPP systems that are homologous to the NAPHQ and NVPHV systems in *E. faecalis*. We propose an evolutionary model whereby the dual QS system was introduced sequentially, with the NVPHV system being the older system and existing in the common ancestor of the *E. faecalis* group of the enterococcal phylogeny, comprising *E. moraviensis*, *E. haemoperoxidus*, and *E. caccae*. The NAPHQ system is newer and likely arose through a duplication and divergence event that coincides with the split and speciation of E. faecalis from the rest of the enterococcal species in the group.

Out of all open reading frames annotated in 1,661 HMP reference genomes, we detected 6,239,015 proteins of all sizes. Of these, 138,792 were smORFs (≤ 50 amino acids) with a start and stop codon. Among these smORFs, 21,860 were predicted to be secreted based on analysis with Phobius, which predicts signal peptide sequences that direct proteins for secretion. By analyzing a 1-gene window upstream and downstream of the putative secreted smORFs (n = 43,720), we identified 342 small secreted proteins with an adjacent TPR-containing protein. After removing redundancies (i.e. multiple instances of the same pair of genes occurring in the same species), we identified a total of 125 unique putative smORF-RRNPP receptor pairs of genes (Figure 1b; Supplementary Table 1). These systems were distributed across 75 phylogenetically-diverse Gram-positive and Gram-negative bacteria (Supplementary Figure 1) and 1 archaeon (*Methanobrevibacter smithii*).

Among these, one putative communication system, consisting of a putative smORF and RRNPP receptor pair, was found exclusively in the gut bacterium and opportunistic pathogen *E. faecalis*. Further investigation revealed a second putative system highly homologous to the first, located in a different chromosomal location. Both systems were present in every complete publicly available *E. faecalis* genome we inspected (n=948), indicating that these two systems are most likely part of the core genome of *E. faecalis*. Notably, only one other organism, *Streptococcus pyogenes*, is known to have two highly homologous RRNPP systems^35^. In *E. faecalis*, each system contains two genes organized in an antisense orientation in the genome. The first gene encodes a putative RRNPP receptor with an N-terminal DNA-binding domain and 3 TPR domains (Figure 1c); the predicted 3-dimensional structures of the receptors for these two systems are highly homologous (Figure 1d). The second gene encodes a 20-amino acid microprotein. Based on signal sequence prediction tools, the small proteins encode a 15-amino acid signal peptide sequence that directs the peptides for secretion, leaving a pentapeptide, NAPHQ and NVPHV (N→C), as the putative signaling molecules of these communication systems (Figure 1e). The receptors sequences are ∼70% identical at the nucleotide and amino acid levels, as are the small proteins. Thus, a computational approach for identifying new RRNPP-type signaling systems revealed the first chromosomally encoded system in *E. faecalis,* and only the second example of an organism encoding two highly homologous RRNPP systems.

### Evolutionary analysis reveals that the presence of the NAPHQ- and NVPHV Systems correlates with the *E. faecalis* speciation event

In recent years, extensive computational and experimental work has shed light on the evolutionary history and phylogeny of enterococci^33^. All enterococcal species appear to fall into one of four main clades, based on a core-genome SNP-based analysis. The two new RRNPP systems we discovered in *E. faecalis* are highly conserved in the species; we next sought to determine whether these systems are present in closely related organisms, and thus are evolutionarily conserved. To assess this, we attempted to determine whether other enterococcal clades encode one or both of the new RRNPP systems. To this end, we annotated putative RRNPP-like communication systems in reference genomes representing 28 *Enterococcus* and outgroup species as described above and searched for systems that are homologous to either QS system. Nearly all the species in our analysis contained at least one putative RRNPP-like communication system (with the exception of *Tetragenococcus halophilus* and *Enterococcus faecium* - Supplementary Table 2). However, only the *E. faecalis* lineage (Figure 1f) contained RRNPP-like systems that were similar to the NAPHQ- and NVPHV-systems. These species include *E. moraviensis*, *E. haemoperoxidus*, and *E. caccae*. Interestingly, of these organisms in the *E. faecalis* lineage, only *E. faecalis* has both a NAPHQ system and a NVPHV system; the other species only contained one system with NTGHV as the putative signaling micropeptide.

Furthermore, the broader genomic neighborhood of the NVPHV system in *E. faecalis* shares several conserved features among all of the organisms that contain the NTGHV system in *E. moraviensis*, *E. haemoperoxidus*, and *E. caccae* (Supplementary Figure 2). These observations support a model whereby the NVPHV-like system was introduced into enterococci prior to the evolution and diversification of the *E. faecalis* group and the NAPHQ system likely emerged in *E. faecalis* through a duplication and divergence event at the point when it split from the other species in the lineage. Taken together, our analysis suggests that this particular type of communication system was first acquired in the common ancestor to the *E. faecalis* lineage within the Enterococci phylogeny, and then through a duplication and divergence event that coincided with the *E. faecalis* speciation event, the second system emerged.

### The NAPHQ and NVPHV system receptors and microproteins are transcribed and translated

Given the *in silico* discovery of these putative communication systems, we sought to determine whether the receptor and microproteins that were annotated were transcribed and translated. First, we measured the transcription of the both receptors and microprotein signals by RT-qPCR in *E. faecalis* OG1RF cultures grown to saturation in rich (BHI) and defined (DM) media. This RNA-level evidence demonstrated that all four genes were transcribed, and that the microprotein genes were more highly expressed in defined vs. rich media (NAPHQ microprotein - 3-fold difference (*P*=1.2 x 10^-4^); NVPHV microprotein-5-fold difference (*P*=7.2 x 10^-8^); Figure 2a). By contrast, the expression of the receptor genes for both systems were comparable across media conditions (Figure 2a). These results suggest that the RRNPP receptors are constitutively expressed, while the expression of the signaling peptides varies depending on environmental conditions (e.g. nutrient limitation). Furthermore, we found that expression of the smORFs is cell density dependent, suggesting that there is a feed-forward loop in their production and supporting the model that these communication systems are quorum-dependent (Supplementary Figure 3).

**Figure 2.**
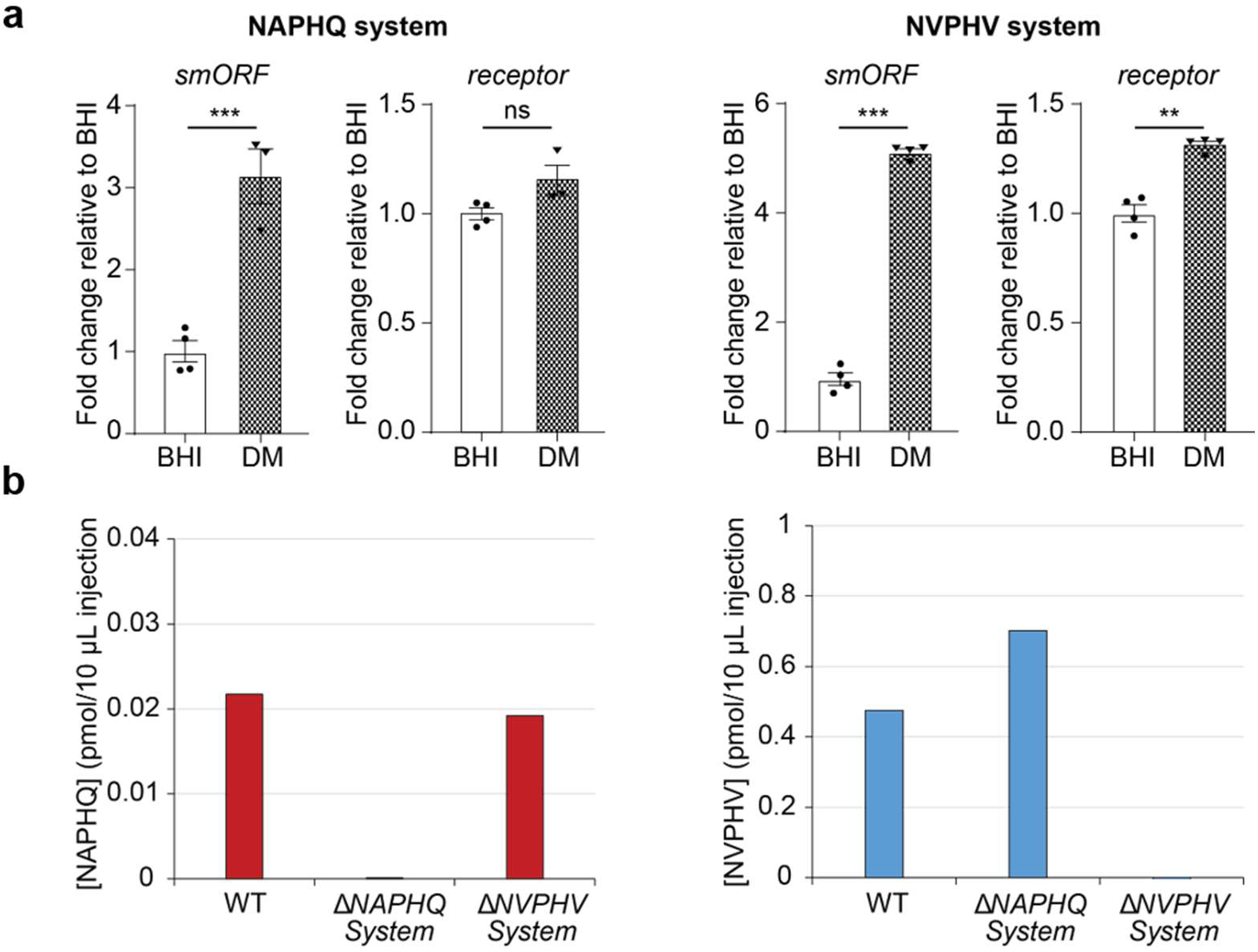
The *E. faecalis* communication systems are transcribed and translated. **a) Expression of the communication systems.** RT-qPCR data is presented demonstrating the relative gene expression levels of the smORF and receptor genes in cultures of *E. faecalis* grown to saturation in either BHI medium (BHI) or defined medium (DM). The expression levels are represented as fold-changes in target gene expression in DM relative to it in BHI. Data are expressed as the mean of n = 4 biological replicates ± standard deviation. Comparisons between groups are performed using a Student’s t test. * P<0.05, ** P<0.01, *** P<0.001. **b) Translational evidence of the signaling peptides.** The two candidate signaling peptides, NAPHQ and NVPHV, were detected by targeted LC-MS/MS analysis from culture supernatants of WT and communication system deletion strains of *E. faecalis*.

We next sought to determine whether the microproteins were translated and if our prediction regarding the sequences of the mature, processed gene products was accurate. To do so, we performed liquid chromatography tandem mass spectrometry (LC-MS/MS) of enriched culture supernatants. As the microproteins are predicted to contain a signal peptide, which is presumably cleaved by the Sec-associated signal peptidase or extracellular proteases, the mature peptides that we expected to detect are pentapeptides (NAPHQ and NVPHV) (Figure 1a, 1e). For this purpose, the strains were grown to saturation in defined media, the culture supernatant was clarified by centrifugation, filtered, subjected to solid-phase extraction to enrich the samples for small peptides, and then analyzed by LC-MS/MS. Both predicted pentapeptides were detected; further, they were quantified based on standard curves of synthesized peptide standards (Figure 2b). As a control, we analyzed the culture supernatant of deletion mutants of the QS systems for the presence of these pentapeptides. Specifically, strains with targeted deletion of both components (receptor and microprotein) of the NAPHQ system and NVPHV system were made, as was a double mutant. These mutants were verified using whole genome sequencing (Supplementary Data). As expected, NAPHQ was absent from the culture supernatant of a deletion mutant of the NAPHQ system, and conversely NVPHV was undetectable in the culture supernatant of a deletion mutant of NVPHV system (Figure 2b). In summary, we find that both QS peptides are transcribed and translated, and we confirm that they are secreted extracellularly as per the canonical model of RRNPP QS systems.

### The NAPHQ and NVPHV systems control *de novo* purine biosynthesis

Having demonstrated that these systems are transcribed and translated, we sought to determine what genetic programs these systems control. Most peptide-based QS systems in Gram-positive organisms control the transcription of downstream programs through the binding of specific promoter sequences within the genome. We therefore hypothesized that the predicted ‘receptors’ function as transcriptional regulators upon binding to their cognate QS peptide signals. To evaluate the effect of the QS peptides on gene regulation in *E. faecalis* and to identify the putative transcriptional targets of these QS systems, we grew wild type *E. faecalis* (HM201) in defined media to log-phase and treated with 5 µM of synthetic pentapeptides (NAPHQ, NVPHV, or their corresponding scrambled controls, which were used as negative controls) for 15 minutes. Treatment of *E. faecalis* with NAPHQ led to the increase in the expression of all 10 genes involved in *de novo* purine biosynthesis along with 2 other genes (the guanine/hypoxanthine permease *pbuO* and the nucleobase transporter *PlUacP*) involved in the transport of purine nucleobases (Figure 3a-3b, Supplementary Table 3, 4). Conversely, treatment of *E. faecalis* with NVPHV resulted in the decrease in the expression of *de novo* purine biosynthesis genes (Figure 3b). *De novo* purine biosynthesis is the process by which bacteria produce purine nucleotides, the building blocks of DNA and RNA from intermediates of central metabolic pathways. The pathway produces inosine 5’-monophosphate (IMP), the precursor of the purine nucleotides adenosine monophosphate (AMP) and guanosine monophosphate (GMP) (Figure 3b). The genes involved in this pathway are organized together in a single polycistronic operon (Figure 3c). We note that when we exposed *E. faecalis* to the full length smORFs, we did not observe significant changes in gene expressions, suggesting that the mature pentapeptides are required to elicit the transcriptional response, and that cleavage and maturation of the signaling peptides does not occur in the extracellular environment (Supplementary Figure 4).

**Figure 3.**
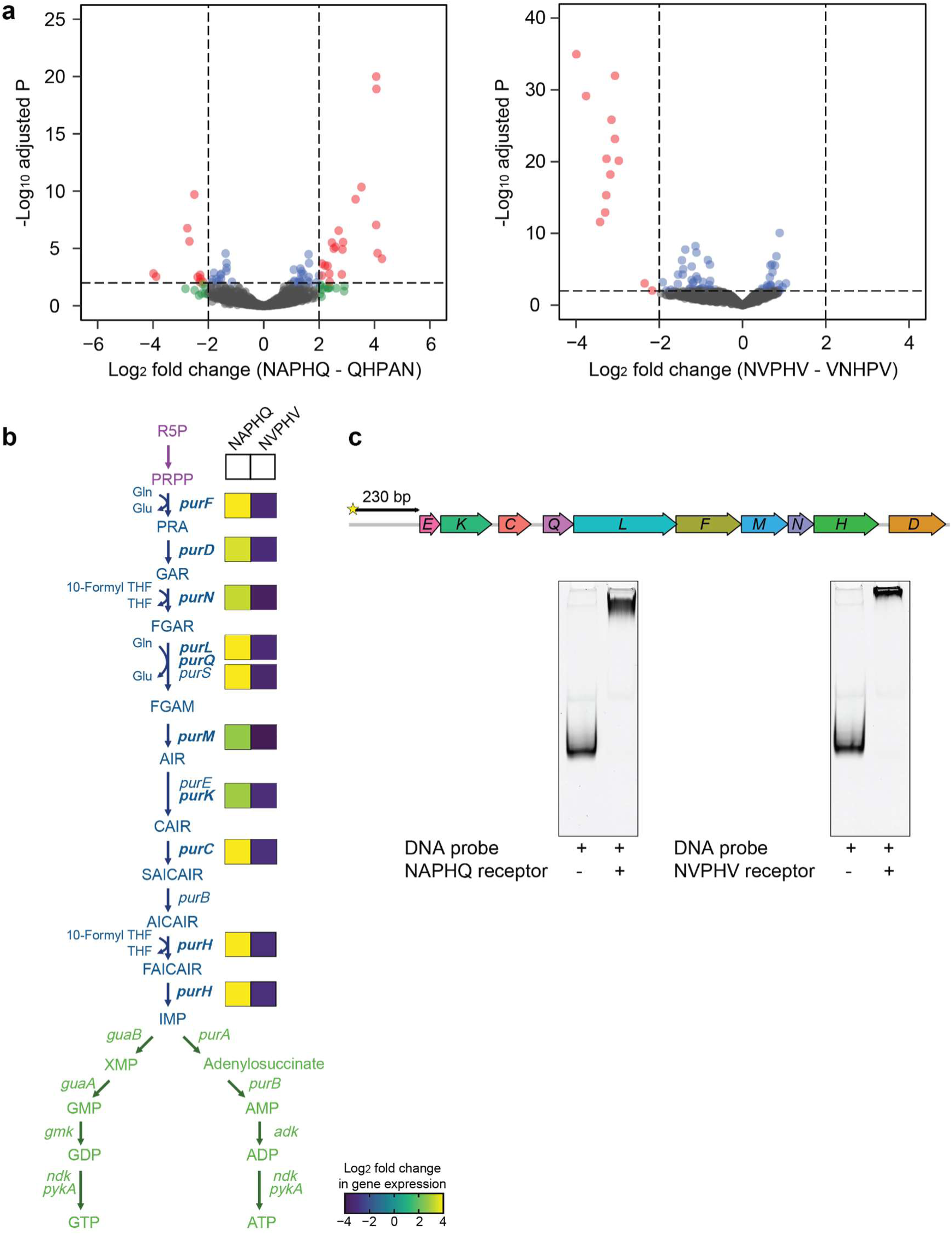
The QS systems modulate *de novo* purine biosynthesis in *E. faecalis*. **a) The signaling peptides elicit a distinct transcriptional response.** Volcano plots demonstrate gene expression analysis of *E. faecalis* grown in DM and treated with 5 μM of the peptides NAPHQ or NVPHV or their respective scrambled controls for 15 min. Differentially expressed genes are defined as those that display at least 4-fold change in gene expression relative to the scrambled control with FDR cutoff < 0.01. **b) Effect of signaling peptides on de novo purine biosynthesis.** A schematic diagram of *de novo* and salvage pathways of purine biosynthesis in *E. faecalis* is shown. *De novo* purine biosynthesis (blue) in bacteria begins with the molecule 5-phosphoribosyl-1-pyrophosphate (PRPP), which is derived from the pentose phosphate pathway intermediate ribose-5-phosphate (R5P). PRPP then proceeds through a series of enzymatic reactions to eventually produce inosine 5’-monophosphate (IMP), a precursor to the purine nucleotides adenosine monophosphate (AMP) and guanosine monophosphate (GMP). Enzymes within the salvage pathway (green) are responsible for making purine nucleoside di-and triphosphates as well as recycling purine bases and nucleotides available in the environment. Treatment of *E. faecalis* with 5 μM of the peptides NAPHQ or NVPHV for 15 minutes results in changes in the transcription level of the genes involved in *de novo* IMP biosynthesis. The log_2_-fold change in gene expression relative to the corresponding scrambled controls is shown in the boxes next to the genes along the pathway. **c) EMSA detection for direct binding of the smORF receptors to the *pur* promoter region.** Agarose gel showing the electrophoretic mobility shift assay (EMSA) using purified recombinant QS system receptors and a 5′-FAM-labeled DNA probe corresponding to the 230 bp region promoter region upstream of the start codon of *purE*, the first gene in the *pur* operon. When the DNA probe is incubated with either QS receptor, the complex migrates at a slower rate through the gel relative to the free DNA probe.

Based on these results, we predicted that the receptors in both systems likely bind to the promoter sequences of the genes that drive purine biosynthesis. The regulation of purine biosynthesis has been extensively studied in other bacteria, such as *Lactobacillus lactis*, *Bacillus subtilis*, and *Staphylococcus aureus*. To assess whether the transcriptional signature of the QS systems was mediated by a direct interaction between the receptors of the QS systems and the *pur* operon promoter region, we carried out an electrophoretic mobility shift assay (EMSA) using purified recombinant QS system receptors and a fluorescent DNA probe corresponding to the 230 bp region promoter region upstream of the start codon of *purE*, the first gene in the *pur* operon. This region contains a predicted *pur* box. Upon incubating the DNA probe with either QS receptor, the complex migrates more slowly in the gel matrix during electrophoresis when compared with the free probe (Figure 3c). Taken together, these results suggest that the both receptors of the QS systems act as transcription factors that directly interact with the *pur* operon, presumably through their DNA-binding domain, and modulate *de novo* purine biosynthesis in *E. faecalis*. Taking together the predicted role of these systems in regulating purine biosynthesis and the order in which they emerged evolutionarily, we named the smORFs encoding genes for NVPHV and NAPHQ *pqs1* and *pqs2* (<u>p</u>urine <u>q</u>uorum <u>s</u>ignal), respectively, and we name their corresponding receptors *pqr1* and *pqr2* (<u>p</u>urine <u>q</u>uorum receptor).

### The *pqs2-pqr2* system is required for robust growth in purine-limited media

Given that we successfully created single and double deletion mutants of each of the QS systems, we sought to determine whether the deletion of either system impacts the optimal growth of *E. faecalis in vitro*. To this end, we grew the WT, single, and double deletion mutants of the *pqs1-pqr1* and *pqs2-pqr2* systems in defined media without purines to saturation and performed a spot dilution assay on selective plates (Figure 4a). We observed that the Δ*pqs2-pqr2* mutant displayed a growth defect relative to the WT strain. Interestingly, the Δ*pqs1-pqr1* mutant showed no such defect; deleting both systems retained the same growth phenotype as in Δ*pqs2-pqr*2, suggesting that there was no additive effect from the loss of both systems. Thus, while these systems are not required for viability *in vitro*, the Δ*pqs2-pqr2* and double deletion mutants exhibit compromised growth.

**Figure 4.**
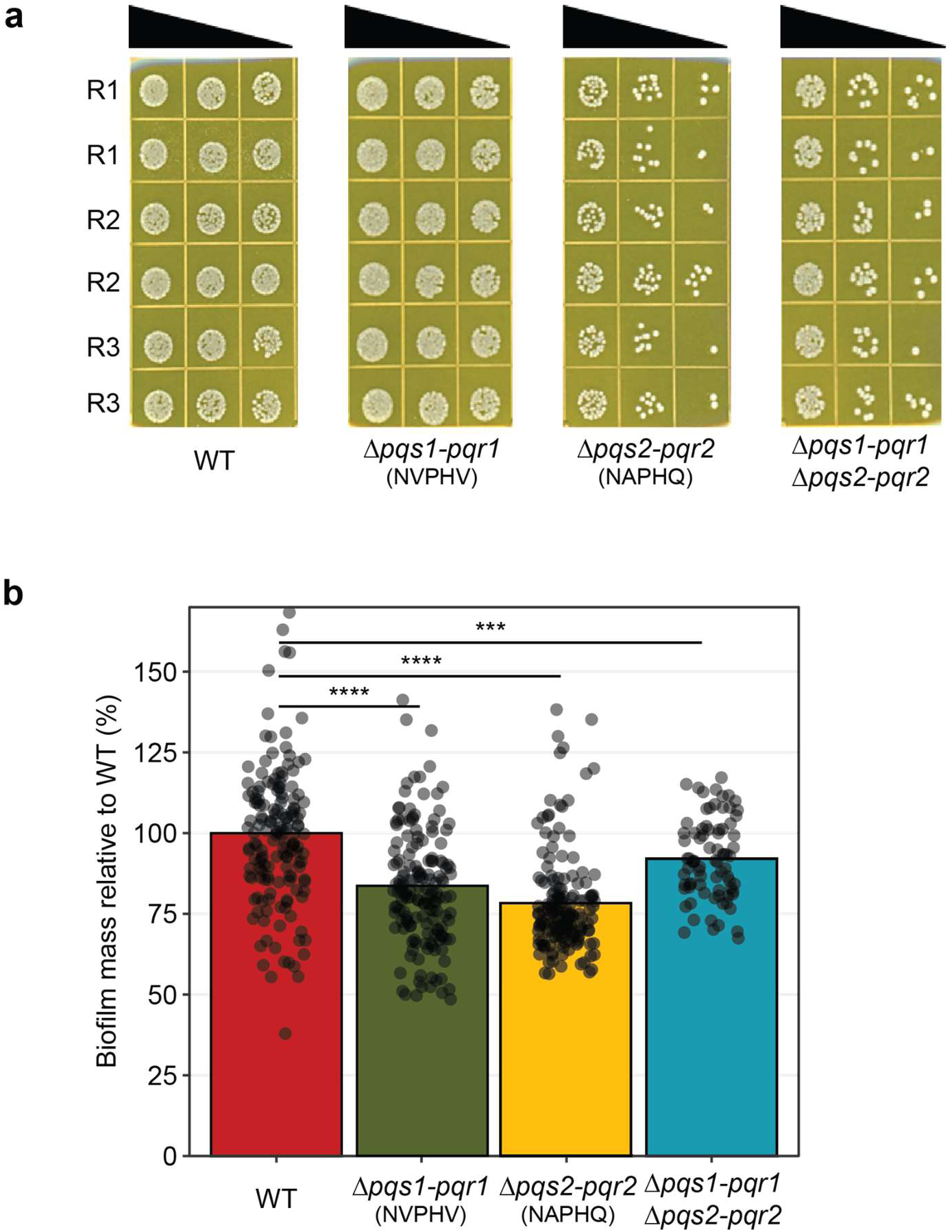
Deletion of QS systems impacts *in vitro* phenotypes of *E. faecalis*. **a) The *pqs2-pqr2* system is required for robust growth in defined media without purines.** Spot dilution assay of E faecalis WT and KO strains (n = 3 biological replicates, each spotted in 2 technical replicates) grown in defined media at 37°C for 18 hours and 10-fold serially diluted and spotted on BHI-RIF plates. **b) Mutants in the signaling systems are defective in biofilm formation.** The mass of biofilms formed by *E. faecalis* wild-type and deletion mutants in the signaling systems was quantified by crystal violet staining. Strains were grown in DMYE in 96-well microtiter plates for ∼18 hours at 37°C without shaking to allow for the biofilms to grow. After multiple washes, biofilms were solubilized in ethanol:acetone (80:20 v/v). Absorbance was measured at 570 nm. The biofilm mass for each strain is expressed relative to its mass in the wild-type strain.

### The QS systems are critical for maximal biofilm production in *E. faecalis*

*De novo* purine biosynthesis is the most highly upregulated pathway in biofilms of *E. faecalis* and *S. aureus* compared to planktonic growth^43^. Given this, a communication system that enables robust purine biosynthesis might be advantageous in the context of a biofilm. To assess this, we tested whether deletion of the QS systems impacts biofilm production in *E. faecalis* and found that there is an up to ∼25% decrease in the biomass of biofilm produced by the single and double deletion mutants in the QS systems relative to the WT strain (Figure 4b). These results indicate that while the QS systems are not essential for biofilm formation in *E. faecalis*, they are important for maximal biofilm production.

### The QS systems are important for intracellular survival of *E. faecalis* within human macrophages

A distinguishing feature of *E. faecalis*, compared to many other pathobionts, is its ability to survive inside macrophages. This ability has contributed to *E. faecalis’* success and versatility as a pathogen. Several groups have shown that *E. faecalis* can sustain robust growth within macrophages for up to 72 hours post-infection, outperforming several other bacterial species, including the closely-related *L. lactis*^44^. Several obligate and facultative intracellular pathogens have a strict requirement for purine biosynthesis for their survival and replication, either due to the lack of host purine nucleotides or because *de novo* synthesis is favored over the import of purines in the intracellular environment^45,46^. Given that survival of *E. faecalis* within macrophages is thought to be critical for its pathogenicity, we postulated that the purine biosynthesis compromised Δ*pqs1-pqr1* and Δ*pqs2-pqr2* mutants would have lower survival in macrophages. To test this, we measured the survival of the WT, single, and double deletion mutants in *pqs1-pqr1* and *pqs2-pqr2* in a human macrophage cell line (Figure 5a). Briefly, we infected U937 human macrophage cells with either the WT, single, or double QS deletion mutants. After 24 hours, the macrophages were lysed, and the recovered bacteria plated on selective plates. Based on these counts, the QS deletion mutants were significantly impaired in their survival within macrophages relative to the WT strain. These results show that the QS systems are important for optimal intracellular survival within macrophages.

**Figure 5.**
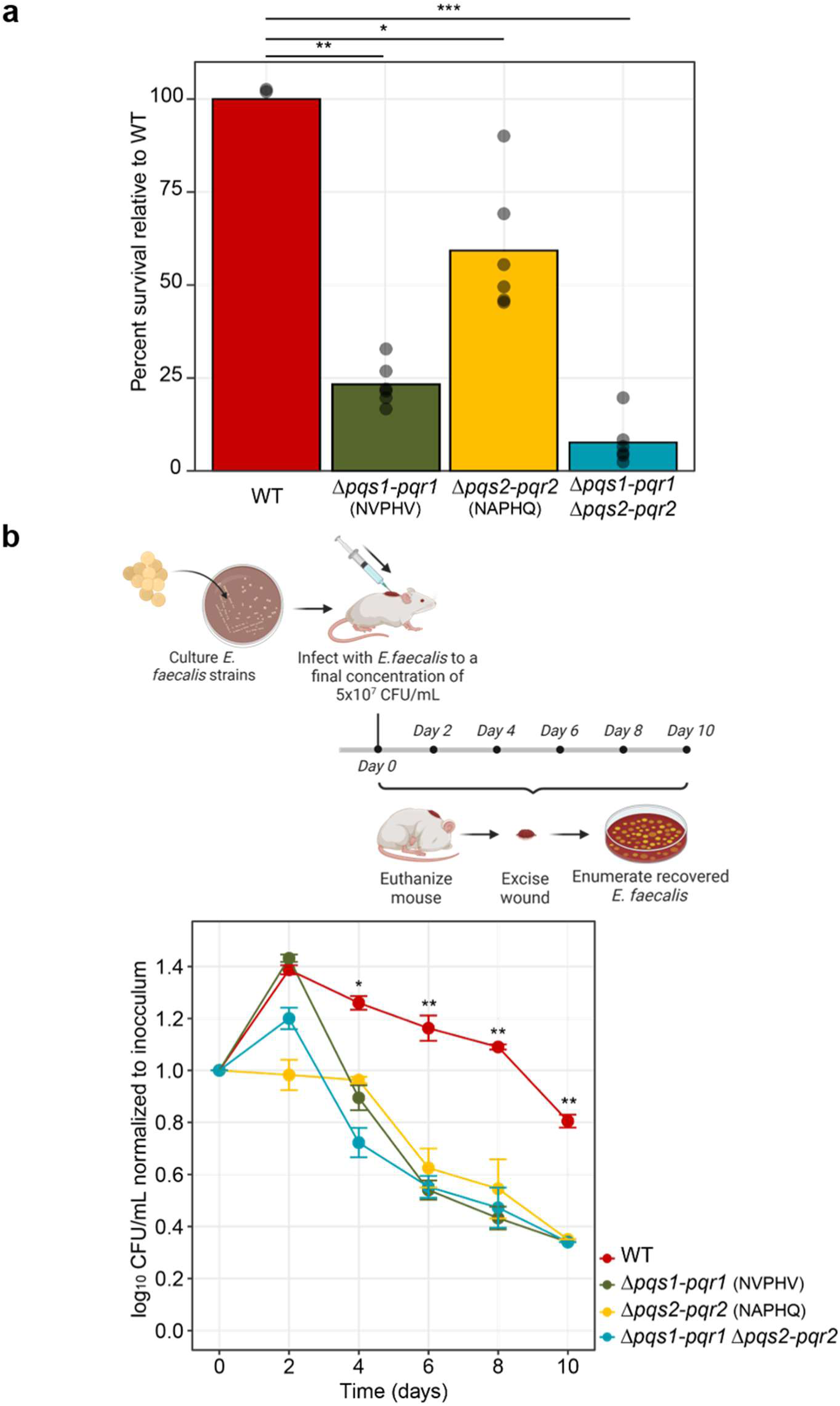
The QS systems are important for optimal *E. faecalis* fitness at the host interface. **a) Fitness of mutants in the signaling systems is impaired inside human macrophages.** U937 human macrophage cell line was infected with E. faecalis wild-type or mutant strains (n = 6 wells per strain). At time intervals, the macrophages were lysed and the recovered bacteria plated. Based on the colony-forming unit (CFU) counts of the recovered bacteria, the % survival of mutants within macrophages is expressed relative to the survival of the WT strain 24 hours post-infection. Comparisons between groups are performed using a Student’s t test. * P<0.05 and ** P<0.01. **b) Mutants in the smORF systems are impaired in growth *in vivo*.** Skin abscess infection mouse model with *E. faecalis* OG1RF wild-type and signaling systems knock-out strains. The back of each mouse was sterilized and shaved, and a superficial skin abrasion was made with a needle prior to application of 20 µL of bacteria in saline solution (at 5×10^7^ CFU per mL) to the abraded area. The progress of infection was monitored every two days for 10 days. At time intervals, animals (n = 4 mice per group) were euthanized and the area of scarified skin was excised, homogenized, and the CFU’s were quantified by 10-fold serial dilution of the homogenate on selective plates. Comparisons between groups are performed using a Student’s t test. * P<0.05 and ** P<0.01.

### The QS systems are important for *E. faecalis* fitness in a murine *in vivo* infection model

Given that the QS systems appear to be important for *E. faecalis* fitness in host-facing phenotypes, such as intracellular survival, we sought to determine whether these systems are important in a relevant organismal infection model. Specifically, we tested the viability of the QS mutant strains in a murine skin abscess infection model. Briefly, a superficial skin abrasion was made on the back of a mouse, and this lesion was infected with a fixed inoculum of either the WT or the QS single and double deletion mutant strains. At 2-day intervals over a period of 10 days, the mice were euthanized, and the area of scarified skin was excised, homogenized, and the recovered bacteria were enumerated by serial dilution plating (Figure 5b). We observed an initial increase in the bacterial load in the wounds infected with the Δ*pqs1-pqr1* mutant or the double QS deletion mutants and a relative stability in the bacterial load of the Δ*pqs2-pqr2* mutant 2 days post-infection. Subsequently, the bacterial counts for each of the single or double deletion mutants dropped substantially relative to the WT strain. In summary, the QS systems are essential for the fitness of *E. faecalis* in the host.

## DISCUSSION

Bacterial peptide-based communication systems are important in *cis*-regulation of various functions in bacterial populations. However, only a few such systems are known to date^25^, likely due to technical challenges in the annotation of microproteins in microbial genomes. Here, by leveraging recent and improved computational approaches for annotating microprotein genes, we identified hundreds of novel candidate peptide-based communication systems in human associated bacterial genomes. These putative systems are distributed across phylogenetically diverse bacteria, including non-Firmicutes. With the development of new tools to annotate RRNPP systems^47^, it would be interesting to further explore the full extent of the phylogenetic diversity of this type of peptide-based communication system.

Among the systems we identified, a particularly interesting set of two systems (which we call *pqs1/pqr1* and *pqs2/pqr2*) were identified in the gut ‘pathobiont’ *E. faecalis.* Using a combination of comparative genomics, microbiological, transcriptomics, targeted proteomics and animal model experiments, we discovered that these two peptide-based QS systems regulate *de novo* purine biosynthesis, the first evidence, to our knowledge, of a QS system regulating central metabolism. Using cell-based and animal model experiments, we further demonstrated that these two systems are important for *E. faecalis* fitness and survival within the host.

Purine accessibility is critical for all bacteria. Some bacteria must scavenge purines from the environment, while others, including many important human pathogens, are capable of synthesizing purines *de novo.* This is important because free host-derived purines are not accessible to extracellular or intracellular bacteria. Thus bacteria must either possess the ability to synthesize purines *de novo* or have the necessary nucleases and transporters to extract purine building blocks from extracellular DNA. In this context, the emergence of *de novo* purine biosynthesis in enterococci marks a critical evolutionary advance for this genus, granting species within the phylogeny the ability to synthesize their own purines. Among the enterococci, *E. faecalis* has been described as the best ‘generalist’ - it can thrive in different host compartments with highly variable concentrations of purines, including the gut, urogenital tract, bloodstream, inside macrophages, and within tissues. We posit that the ability to efficiently fine-tune purine biosynthesis at the population level through the *pqs1/pqr1*-*pqs2/pqr2* QS systems is a key feature in enabling *E. faecalis* to adapt to these different environments.

When viewed through an evolutionary lens, *E. faecalis* likely acquired the ability to regulate purine biosynthesis through QS in two steps. The first of the two systems, *pqs1/pqr1*, appears to be evolutionarily older, as it is found in *E. faecalis* as well as in organisms in a neighboring evolutionary branch (*E. moraviensis*, *E. haemoperoxidus*, and *E. caccae*). While we do not have experimental evidence regarding the function of the *pqs1/pqr1* homolog in organisms of this neighboring branch, it may also be involved in regulating purine biosynthesis. We posit that the second, newer system likely arose through a duplication and divergence event. This second system (*pqs2/pqr2*) is unique to *E. faecalis*, suggesting that it plays a role in differentiating *E. faecalis* from closely related species. We find that while not essential, both systems are critical for the fitness of *E. faecalis*, particularly in its interaction with the host. Our finding that purine biosynthesis is regulated by multiple QS mechanisms suggests that tight regulation of purine levels in *E. faecalis* is important for its lifestyle. The ability of *E. faecalis* to carefully and dynamically regulate purine biosynthesis likely enables it to thrive in highly variable purine-containing niches. Consistent with this model, our data show that these QS systems are important for maximum biofilm production, survival within macrophages, and fitness *in vivo* in a wound infection model. Furthermore, our *in vivo* results are concordant with recent work that identified purine biosynthesis genes as critical for the replication and persistence of *E. faecalis* during wound infection and catheter-associated urinary tract infection^48^.

Curiously, our results present an intriguing paradox: why does *E. faecalis* possess two QS systems that oppositely modulate purine biosynthesis at the transcriptional level, yet have similar impacts on host-facing phenotypes, such as survival within macrophages and survival in a wound infection model? The simplest model we can deduce from these data is that, at the mechanistic level, the two systems are antagonistic to one another -*pqs1-pqr1* acts as a repressor of *de novo* purine biosynthesis, and *pqs2-pqr2* acts as an activator. The *in vivo* phenotypes can be potentially explained by positing that purine levels that are either too high or too low are detrimental to *E. faecalis* in these settings, and thus careful and tight regulation of exact purine levels is critical to optimal survival. While we provide some insight into the coordinated regulation and consequences of these two systems, ongoing and future work will focus on resolving the question of why a dual QS system for *de novo* purine biosynthesis modulation exists and the biochemical model by which the two systems interact and perform their function.

While we have discovered and demonstrated the function of two new QS signaling systems *in vitro* and *in vivo*, our study has several limitations. First, we do not yet fully understand how the two systems interact with each other and whether they display cross-talk or cross-regulation.

Second, we do not know how these QS systems sense purine levels. Beyond their role as energy-carrying compounds and building blocks of nucleic acids, purine metabolites are also signaling molecules that regulate different functions within the cell^49–51^ or mediate microbe-microbe interactions^52^. Recent work has uncovered a widely-distributed purine-binding motif in the sensor domains of thousands of bacterial receptors implicated in various functions^53^. It is intriguing to speculate whether the QS systems discovered in our work interface with other purine-sensing receptors in *E. faecalis*. Third, the interaction between these QS systems and the classical purR-mediated regulation of *de novo* purine biosynthesis or the purine salvage pathway is not yet known. Carefully controlled experiments using metabolomics-based approaches and varying levels of exogenous purines might provide some answers to these mechanistic questions.

This study provides insights into the evolution and function of two new antagonistic peptide-based QS in the biology and lifestyle of one of the most prevalent enterococcal species in the human gut and an important hospital-associated pathogens. We posit that having an additional layer of regulation of purine biosynthesis at the population level has given *E. faecalis* the ability to thrive in highly variable purine-containing niches, making it the most successful generalist among human-associated microbiota. Notably, while *E. faecalis* is classified as a commensal gut bacterium, it can also be pathogenic and, along with other *Enterococcus spp.*, presents a major burden in the clinic due to their intrinsic and acquired antibiotic resistance. Establishing the critical role of these QS systems for *E. faecalis* fitness *in vivo* in a murine infection model validates these systems as promising anti-*E. faecalis* targets. Understanding this communication system will help devise strategies to block it and prevent enterococcal domination, which is associated with *Enterococcus* bacteremia and significant morbidity in hospitalized patients.

## METHODS

### Identification of putative peptide-based communication systems

The Human Microbiome Project reference genomes files (n = 1661 complete genomes as per HMP website) were downloaded from https://www.hmpdacc.org/hmp/HMRGD/#data. Prodigal was used to annotate all open reading frames with adjusted thresholds to detect ORFs as short as 20 amino acids. A total of 6,239,015 proteins of all sizes were detected, 138,792 of which were smORFs (≤ 50 aa) with a start and stop codon. Those predicted ORFs that were 50 residues or fewer (with a lower threshold of 20 residues which is the lowest threshold that can be used to successfully run prodigal annotation on all the genomes.) were then evaluated using Phobius^54^ to identify those with a predicted signal peptide, to determine which peptides were likely to be secreted. TPRpred^55^, which predicts proteins that contain Tetratricopeptide Repeat (TPR) domains, was run on the proteins that are 1 gene upstream or downstream of the putative small secreted protein and those with ≥ 50% probability of containing a TPR domain were scored as positive. Through this analysis, we identified putative peptide-based communication systems, each composed of a small, secreted protein and an adjacent TPR-containing protein. RRNPP-like systems were identified in enterococcal genomes using the same criteria as described above (Supplementary Table 2). The full sequence smORFs were clustered using CD-HIT^56^ with a 60% identity threshold. The representative sequences from the smORF clusters were visually inspected to identify those that looked similar.

### Building a phylogenetic tree of the species with putative RRNPP systems

76 HMP genomes representing the microbial species in which we identified the 125 putative RRNPP systems were used to build a phylogenetic tree. The tree was built with GTDB-tk v2.3.0^57^ de_novo workflow. Briefly, for each genome 120 bacterial marker genes were identified, these marker genes were used to create multiple sequence alignment and a phylogenetic tree was inferred based on the multiple sequence alignment using FastTree. The tree was visualized in iTOL^58^. The phyla were colored based on the GTDB-tk classification workflow with default parameters.

### Culture media

Brain heart infusion (BHI) broth (Millipore-Sigma 53286) was prepared as per product specification. Defined media (DM) was prepared as follows. Per 500 mL of media: 3.5 g of dipotassium hydrogen phosphate (K_2_HPO_4_), 1 g of potassium dihydrogen phosphate (KH_2_PO_4_), 50 mg of magnesium sulfate (MgSO_4_), 7.5 g of D-glucose (1.5% w/v final concentration), mineral supplement solution (ATCC MDMS) and vitamin supplement solution (ATCC MDVS) to a final concentration of 1%, and 0.5 g of each of the 20 proteinogenic amino acids. The pH of the media was adjusted to pH 7.2, filter-sterilized through a 0.22 µm filter, and stored at 4 °C. DMYE was prepared by adding yeast extract (Fisher Scientific BP1422-500) to DM to a final concentration of 0.2%.

### Generating deletion mutants in the quorum-sensing systems

Deletion mutants of the peptide quorum sensing systems were generated in *E. faecalis* OG1RF in a 2-step procedure based on homologous recombination as described in ^59–61^. All primers used for making the deletion mutants are in Supplemental Table 5. For each deletion strain, the mutant allele in which the entire quorum sensing system (smORF + receptor) is deleted was constructed. Each of the putative QS systems along with a 1000 bp of flanking DNA from up- and downstream from the of the systems and cloned into the temperature-sensitive vector pJH086 (courtesy of Chris Kristich lab) using Gibson assembly^62^ (NEB). The construct was then transformed into *E. faecalis* by electroporation. Electrocompetent cells were prepared using established protocols ^63,64^. Cells were suspended in 1000 µl BHI and allowed to recover at 30 °C for 1.5-2 hours prior to plating on selective agar (BHI with 10 µg/ml Chloramphenicol and 150 µg/ml X-Gal) at 30°C (permissive for pJH086 replication). Pale blue colonies typically appeared within 24-36 hours. Four to eight pale blue transformants were restreaked 1x on the same selective medium as above for single-colony purification. Single-colony-purified transformants were restreaked on the same selective medium as above at 42 °C (non-permissive temperature). Dark blue colonies in which the pJH086 derivative has integrated into the chromosome typically appeared within 24-36 hours. These integrants were restreaked 2x on the same selective medium at 42 °C for single-colony purification. Colony PCR was used to verify that the pJH086 derivative plasmid integrated into the genome in the correct locus. 2-4 verified single integrants derived from independent transformants were streaked directly on the counterselection medium MM9YEG agar supplemented with 10 mM p-ClPhe and 150 µg/ml XGal at 30°C. Isolated white colonies in which the integrated plasmid has been excised and lost by segregation appeared within ∼24 hours. The colonies were patched on chloramphenicol plates to confirm their sensitivity (due to the excision of the integrated plasmid), then re-streaked on counterselection media to further purify them, and screened by PCR and Sanger sequencing to determine that they carry wild-type or deletion mutant allele. A deletion mutant of each system was created. The double deletion mutant in both systems was created sequentially by introducing the second deletion in competent cells of the first deletion mutant.

For 100 mL of MM9YEG with 10 mM p-Cl-Phe, the following was added: 0.25 g yeast extract, 199.6 mg p-Cl-Phe (Sigma C 6506), 1.6 g Bacto agar (Becton-Dickinson cat# 214010), water up to 89 mL, and a stir bar. Immediately after autoclaving for 30 minutes, the medium was thoroughly mixed to fully dissolve p-Cl-Phe and the mix maintained at 55 °C on a hot plate. Then the following was added: 10 mL of 10x sterile M9 salts (made with 60 g anhydrous Na_2_HPO_4_, 30 g KH_2_PO_4_, 5 g NaCl, 10 g NH_4_Cl per liter and autoclaved for 30 minutes), 0.5 mL sterile 50% glucose, 150 µg/mL XGal, and 20% sucrose. The media was mixed and the plates poured.

### RT-qPCR assays

Overnight cultures of E faecalis HM201 were set up in BHI and grown at 37 °C shaking at 150 rpm. The overnight cultures were diluted 1/100 in either BHI or defined media (DM) at 37°C, shaking at 150 rpm and were grown overnight to saturation. Cultures (6-7 mL) were collected and placed on ice and cells were pelleted at 9,200 x g at 4 °C for 10 minutes. For RNA extractions, cell pellets were resuspended in lysis buffer (250 μL 1X PBS + 5 μL lytic enzyme (Qiagen) per sample) and incubated at 37 °C for 30 minutes. A volume of 30 μLof 20% SDS was added to each sample then the samples were incubated at 37 °C for another 30 minutes. A volume of 1.5 mL Trizol was added to each cell lysate followed by a 10-minute incubation on the bench. A volume of 0.5 mL chloroform was added to each sample, mixed vigorously for 15 seconds, incubated on the bench for 3 minutes, and centrifuged at 13,200 x g for 10 minutes. The aqueous phase (∼800-900 μL of the supernatant) was transferred to a new tube, mixed with an equal volume of isopropanol, and incubated on the bench for 20 minutes. Samples were then spun down at 13,200 x g for 10 minutes at 4 °C. The pellets were washed twice with 75% ethanol then left to air dry on the bench. RNA was eluted with 50 μL of RNAse-free water and samples were stored at -80 C. For RT-qPCR measurements, equal amounts of RNA (100 ng) were set up in 384-well plates using a Biomek FX liquid handler (Applied Biosystems) and processed as follows. Samples were DNAse treated with DNAse I (NEB) in 6 μL reactions.

Reactions were run at 37 °C for 10 minutes followed by heat inactivation at 75 °C for 10 minutes then the reactions were carried over to the cDNA synthesis step. 20 μL reactions were set up as per the AffinityScript Multiple Temperature cDNA Synthesis Kit protocol (Agilent), then diluted with 80 μL of nuclease-free water and stored at -20 °C. For qPCR, 10 μL reactions were set up with 2 μL of each cDNA preparation using the Luna Universal qPCR Master Mix protocol and primers to amplify the genes encoding the small proteins (NAPHQ smORF and NVPHV smORF), their receptors (NAPHQ receptor and NVPHV receptor), *recA* and *rpoZ* as housekeeping genes (Supplementary Table 5). The relative fold gene expression in the samples was calculated using the ΔΔCq method and expressed as fold change in gene expression in DM relative to BHI media.

### Proteomics assays to measure micropeptides

In order to detect and quantify the micropeptides in culture supernatants of *E. faecalis*, overnight cultures of the wild-type and deletion mutant strains were set up in BHI broth with rifampicin (100 μg/mL) at 37 °C, 180 rpm. The cultures were diluted 1/100 in 350 mL of defined media supplemented with 0.2% yeast extract (to promote maximum culture growth) and grown overnight at 37 °C, 180 rpm. The following day, the cultures were centrifuged at 9,200 x g for 20 minutes. The supernatants were then filtered through 0.2 μm filters to remove any bacterial cells, and stored at 4 °C until the next step of processing. To optimize the detection of secreted small peptides, 100 mL of medium alone or 300 mL of filtered bacterial culture supernatants was enriched using C_18_ silica resin, as follows. The samples were acidified with the addition of glacial acetic acid to a final concentration of 0.5 N. The acidified samples were filtered through 0.45 μm filters and purified using Bond Elut C_18_, 40 μm, solid phase extraction cartridges (Agilent, Santa Clara, CA), 1 x 1 g C_18_ for 100 mL medium alone and 4 x 1 g for 300 mL of culture supernatant. Cartridges were primed with methanol and equilibrated with two column volumes of triethylammonium formate (TEAF), pH 3. Samples were applied and cartridges were washed with multiple volumes of TEAF, pH 3 buffer. The peptide enriched fraction was eluted with 75% acetonitrile/25% TEAF pH 3 and evaporated to dryness using a SpeedVac concentrator. Lyophilized samples were reconstituted in 1 mL water and then centrifuged at 15,000 x g for 10 minutes to remove insoluble material. An aliquot was used to measure total protein content and the clarified, peptide enriched fraction was used for LC-MS/MS analysis. For LC-MS/MS quantitation, samples were analyzed on a Dionex Ultimate 3000 LC system (Thermo) coupled to a TSQ Quantiva mass spectrometer (Thermo) fitted with an Accucore C_18+_ column (1.5 µm, 100 x 2.1 mm i.d., Thermo). The following LC solvents were used: solution A, 0.1 % formic acid in water; solution B, 0.1 % formic acid in acetonitrile. The following gradient was utilized: 0 % B for 2 minutes, 0-40 % B in 15 minutes, 40-100 % B in 2 minutes, 100 % B for 1 minutes and re-equilibrate at 0 % B for 7 minutes, for a total run time of 25 minutes at a flow rate of 0.1 mL/min. The injection volume was 10 µL, the column oven temperature was set to 40 °C and the autosampler kept at 4 °C. MS analyses were performed using electrospray ionization in positive ion mode, with spray voltages of 3.5 kV, ion transfer tube temperature of 325°C, and vaporizer temperature of 275°C. Multiple reaction monitoring (MRM) was performed by the following transitions: NAPHQ, 566.3>207.1, 566.3>235.1, 566.3>284.1, 566.3>381.3, 283.7>235.1, 283.7>266.6, 283.7>381.1; NVPHV, 565.4>235.1, 565.4>255.1, 565.4>334.2, 565.4>352.1, 283.3>235.1, 283.3>255.1, 283.3>352.1. Skyline ^65^ was used to measure peak areas, quantitation was obtained by using standard curves of the pure peptides spiked in media.

### Measuring the promoter activity of the *smORF* genes using luciferase reporter assays

To make the promoter-reporter systems, the following DNA fragments were prepared. The *luxABCDE* operon was amplified from pKS310 plasmid (courtesy of Michael Federle lab) with primers SZBP425 and SZBP426. The backbone of the plasmid pGCP123-GFP_g1 (Addgene #153518) was amplified with the primers SZBP427 and SZBP428. The regions upstream of the start codon of the NAPHQ smORF (198 bp) or the NVPHV smORF (197 bp) were amplified by SZBP429 and SZBP430, or SZBP431 and SZBP432, respectively. The fragments were assembled by Gibson assembly^62^ (NEB) to generate promoter-reporter constructs where *luxABCDE* expression is under the control of each P*_smORF_* promoter.

To monitor the activity of the P*_smORF_* promoters *in vitro*, a kinetic assay was adapted from^66,67^ as described below. Reporter constructs (P*_NAPHQ-luxABCDE_* or P*_NVPHV-luxABCDE_*) in *E. faecalis* OG1RF were grown in BHI with kanamycin (500 µg/mL) at 37 °C, 180 rpm. The overnight cultures (4 biological replicates) were diluted 100-fold into different media (BHI, defined media supplemented with 0.2% yeast extract (DMYE), or defined media (DM)) and set up in a Greiner clear-bottom opaque white 96-well plate (cat. no. 655098) (200 µL per well). The inter-well spaces were filled with 1% decanal in mineral oil (50 µL per space). While the *luxABCDE* operon generates its own substrate for luminescence, decanal vapor helps to saturate the assay as it provides an exogenous substrate for LuxA-B to produce luminescence. The plate was placed in a SpectraMax Paradigm Multi-Mode microplate reader (Molecular Devices) and set to incubate at 37 °C. The lid was kept on the plate and sealed with parafilm throughout the kinetic run in the plate reader. At 30-minute intervals, absorbance at 600 nm and luminescence were measured. Relative luminescence was calculated by normalizing luminescence to absorbance values at each timepoint.

### Transcriptomics

All peptide exposure experiments were conducted in 3 biological replicates. Overnights of *E. faecalis* HM201 were set up in BHI and grown at 37 °C shaking at 150 rpm. The overnights were diluted 1/100 in defined media at 37 °C, 150 rpm to OD_600_ ∼ 0.4 (3-4 hours). The cultures were exposed to either the test peptide or its corresponding scrambled control to a final concentration of 5 μM for 15 minutes at 37 °C, 150 rpm. Cultures (4 mL) were then quenched with 0.5 mL of ice-cold quenching solution (90% vol/vol ethanol + saturated acidic phenol). The quenched cultures were incubated on ice for 10 minutes. Cells were pelleted at 9,200 x g at 4 °C for 10 minutes. For RNA extractions, cell pellets were resuspended in lysis buffer (250 μL 1X PBS + 5 μL lytic enzyme (Qiagen) per sample) and incubated at 37 °C for 30 minutes. A volume of 30 μL of 20% SDS was added to each sample then the samples were incubated at 37°C for another 30 minutes. A volume of 1.5 mL Trizol was added to each cell lysate followed by a 10 minute incubation on the bench. A volume of 0.5 mL chloroform was added to each sample, mixed vigorously for 15 seconds, incubated on the bench for 3 minutes, and centrifuged at 13,200 x g for 10 minutes. The aqueous phase (∼ 800-900 μL of the supernatant) was transferred to a new tube mixed with an equal volume of absolute ethanol. RNA was then extracted from the samples as described in the Zymo RNA Clean & Concentrator-5 kit protocol. Column purification was found to be superior in obtaining RNA samples of sufficient quality to be used for RNA sequencing. RNA was eluted from the columns with 25-30 μL of RNAse-free water. RNA extracts were stored at -80 °C. Ribosomal RNA was depleted with the Illumina Ribo-Zero Plus rRNA Depletion Kit (Bacteria) according to the manufacturer’s instructions. cDNA sequencing libraries were prepared with the Truseq Stranded mRNA kit following the Truseq Stranded mRNA LT protocol. Libraries were sequenced with 2 × 150 bp reads on an Illumina NovaSeq 6000 Sequencing System (Novogene), each library receiving ≥ 2 Gb sequence coverage.

Transcriptomics reads (150 bp in length) were quality filtered using trim galore^68^, using default parameters and a quality score cutoff of 30. Reads were mapped to the Enterococcus faecalis strain HM201 reference genome using bowtie2^69^ using default parameters except allowing for no mismatches. For each ORF, mapped reads were counted using bedtools coverage^70^. Differential expression analysis was done using DESeq2^71^. All raw sequencing data will be released on SRA upon publication.

### Cloning, expression and purification of recombinant peptide receptors

To isolate receptor recombinant proteins, constructs were created to overexpress each protein with an C-terminal 6-histidine tag. Briefly, the genes encoding NAPHQ receptor and NVPHV receptor were amplified from *E. faecalis* genomic DNA using Q5 polymerase using the following primers which were designed to contain a ribosomal binding site and a C-terminal 6-histidine tag: for NAPHQ receptor: 5′-GGGGACAAGTTTGTACAAAAAAGCAGGCTTAACTTTAAGAAGGAGATATACATATGAGAGTAGCGGGAG-3′ and 5′-GGGGACCACTTTGTACAAGAAAGCTGGGTTTTAGTGATGGTGATGGTGATGTTTAAAACTGATATTAAACTCT-3′; for NVPHV receptor:5′-GGGGACAAGTTTGTACAAAAAAGCAGGCTTAACTTTAAGAAGGAGATATACATATGAATTTACATAATAATACAAGTGGAG-3′ and 5′-GGGGACCACTTTGTACAAGAAAGCTGGGTTTTAGTGATGGTGATGGTGATGATAGGAAATATTAAAAGC-3′. The PCR products were purified and cloned into pDEST14 using the Gateway cloning and Expression Kit (Invitrogen) as per manufacturer instructions, and the constructs were confirmed by DNA sequence analysis. Each construct was transformed into *E. coli* BL21-AI chemically-competent cells before protein expression and purification. The following procedure was followed for the expression and purification of each of the two proteins. For protein expression, each clone was grown in 1 L of LB with ampicillin (100 μg/ml) at 37 °C with shaking at 200 rpm until the culture reached an OD_600_ ∼ 0.5. The culture was then induced with 0.2% L-arabinose and grown for an additional 4 hours before harvesting by centrifugation at 10,000 x g for 20 minutes. The cell pellets were resuspended and washed with a 0.85% saline solution, pelleted and stored at −20°C. For protein purification, the cell pellets were thawed on ice and resuspended in 50 mL of lysis buffer (HEPES 25 mM, NaCl 300 mM, DTT 5 mM, 30% glycerol, 20 mM imidazole, 0.1% Triton X-100, 100 μg/mL lysozyme, 0.5 mg DNase, 0.5 mg RNase, protease inhibitor cocktail (Roche) pH 7.5). Cells were lysed using a microtip attached to an ultrasonic sonicator (Model 705 - Fisher) on ice over 5 rounds each lasting 30 seconds at amplitude 60. The lysate was clarified by centrifugation in a Sorvall Lynx 4000 Centrifuge at 12,000 rpm for 1 hour at 4 °C. The clarified lysate (∼ 45 mL) was mixed with 3 mL of HisPur Ni-NTA Resin (Thermofisher) and placed on a rocking mixer for 1.5 hour at 4 °C to allow the recombinant protein to bind to the nickel resin. Purification was done by gravity flow by placing clarified lysate nickel resin mix in the column and allowing it to flow through. The resin (∼ 3 mL) was then washed with 45 mL of Buffer A (HEPES 25 mM, NaCl 300 mM, DTT 5 mM, 30% glycerol, 100 mM imidazole, pH 7.5). Recombinant protein was eluted with 6 mL of Buffer B (HEPES 25 mM, NaCl 300 mM, DTT 5 mM, 30% glycerol, 500 mM imidazole, pH 7.5). Fractions were analyzed by SDS-PAGE tricine gels (Supplementary figure 5), and those containing pure His-tagged protein were pooled and buffer exchanged using 10 mL ZEBA desalting columns (Thermofisher) as per manufacturer instructions. The fractions were first buffer exchanged against HEPES 25 mM, NaCl 300 mM, DTT 5 mM, 30% glycerol, 50 mM EDTA, pH 7.5 to chelate any free nickel in the eluate and avoid protein precipitation, then against the final storage buffer HEPES 25 mM, NaCl 300 mM, DTT 5 mM, 30% glycerol, pH 7.5. All centrifugation steps were done at 1000 x g at 4 C. About 10-12 mg were obtained per 1 L pellet for each of the two recombinant proteins. Aliquots of pure protein were stored at −80 °C.

### EMSA assays

The probe DNA fragment corresponding to 230 bp upstream of the start codon of the pur operon in *E. faecalis* was amplified from genomic DNA by PCR using primers containing a 5′-FAM fluorescent tag on the forward primer. The resulting DNA fragments were run on a 4% agarose gel, the bands excised, and gel purified using the Qiagen gel purification kit. The purified probe was eluted in nuclease-free water and stored in opaque tubes at -20 °C. EMSA reactions (20 uL) were set up in the following reaction buffer: 20 mM HEPES, pH 7.9, 100 mM KCl, 12.5 mM MgCl2, 0.2 mM EDTA, pH 8.0, 0.5 mM DTT, 50 µg/mL salmon sperm DNA, 0.001 U/µL poly(dI•dC), 100 µg/mL BSA, 0.5 mM CaCl2, and 12% (v/v) glycerol. Recombinant receptor proteins (4 uM) were added to the reaction mix for 30 minutes prior to the addition of the DNA probe to allow the recombinant protein to interact with the excess non-specific DNA in the reaction mix. Reactions were initiated by adding 10 nM of fluorescent DNA probe. The binding assay was run for 15 minutes at room temperature. To visualize DNA probe migration, a 6% DNA retardation gel (Thermofisher) was pre-run at 90 V for 5 minutes with 0.5X TBE buffer (prepared from 5X TBE buffer (Thermofisher)). Then 8 μL of each sample was loaded directly in the wells of the gel, then the gel was run in the same buffer at 90 V at 4 C. The gel was imaged using the Typhoon biomolecular imager (Cytiva) using the FAM settings (Ex 495 nm, Em 520 nm).

### Spot dilution assays

Overnights of the strains were set up in BHI media with rifampicin (100 μg/mL) and grown overnight at 37 °C at 180 rpm. Cultures were diluted back 1/100 in DM (which does not contain any purines). The subcultures were set up in a sterile clear polystyrene flat-bottom 96-well plate (200 μL per well). The plate was incubated overnight at 37 °C. The cultures were then 10-fold serially diluted and spotted on selective plates (BHI agar with rifampicin at 100 μg/mL). The assay was set up in 3 biological replicates, each spotted in 2 technical replicates.

### Biofilm assay

Overnight cultures of strains of interest were set up in BHI media with rifampicin (100 μg/mL) and grown overnight at 37 °C at 180 rpm. Cultures were diluted back 1/100 in DMYE with rifampicin (100 μg/mL). The subcultures were set up in a sterile clear polystyrene flat-bottom 96-well plate (200 μL per well). The two outer columns (16 wells total) were filled with blank media as a sterility and assay baseline control. Given the high variability of the assay, the assay was set up in multiple biological and technical replicates with 40-80 wells per tested strain. The plates were incubated at 37°C (without shaking) for ∼ 18 hours. At the endpoint, the cultures were carefully removed from the assay plates using a multichannel pipettor taking care not to disrupt the brittle and flaky biofilms that formed at the bottom of the plates. Holding the plates at a 45-degree angle and using gentle pipetting, the wells were washed 3 times with deionized water (200 uL/well) without mixing, then allowed to air dry in an inverted position for 30 minutes. Next, the wells were stained with 200 µL of 0.1% crystal violet per well for 30 minutes at room temperature in the dark (cover the plate with a piece of foil). The crystal violet stain was removed from the plates then washed 3 times with deionized water (200 uL/well). The plates were left to dry completely (several hours to overnight). The biofilms were solubilized with 200 µL/well of ethanol:acetone (80:20), mixed well, and allowed to incubate at room temperature for 15 minutes. The solubilized biofilms were quantified by measuring their absorbance at 570 nm in a spectrophotometer plate reader. Absorbance intensity is directly proportional to the mass of biofilm formed.

### Macrophage infection assays

The U937 cell line (CRL-1593.2) was maintained in suspension culture in RPMI-1640 supplemented with 10% (v/v) heat-inactivated fetal calf serum, 2 mM L-glutamine and 100 U/mL penicillin/streptomycin, at 37°C in a humidified atmosphere of 5% CO2. U937 monocytes were seeded in tissue culture media at 250,000 cells/mL. After 24 hours, cells (∼ 500,000 cells/mL) were induced to differentiate with 100 ng/mL phorbol 12-myristate 13-acetate (PMA) (Sigma) and incubated for up to 48 hours until most cells differentiated into adherent spindle-shaped cells. Once differentiated, media were aspirated, cells were washed with cells were treated with buffered saline (DPBS) without calcium and magnesium, then treated the monolayer with TrypLE (ThermoFisher), incubated at 37 °C for 10 minutes until cells detached from the surface. Media was added to the flask to inactivate the reagent and to collect the cells. The cell suspension was centrifuged at 100 x g for 10 minutes in a 15 mL conical tube. After discarding the supernatant, the cell pellet was resuspended in fresh media without PMA or antibiotics, split into 24-well plates at a density of 100,000 cells/well, and incubated overnight at 37 °C. The following day, the cells are adherent and ready for infection. Overnight cultures of the *E. faecalis* wild-type and deletion strains were set up in BHI with rifampicin (100 μg/mL). The overnight cultures were pelleted and washed 3 times in sterile DPBS, then resuspended in sterile DPBS. For each strain, quadruplicate wells of cells were infected at an MOI of 10 based on CFU counts pre-determined for each of the strains. The plates were spun down at 100 x g for 2 minutes to bring the bacterial cells in contact with the cell monolayer. After incubating the cells at 37 °C for 1 hour, the cells were washed 3 times with sterile DPBS and then incubated in media with vancomycin (16 μg/ml) and gentamicin (150 μg/ml) to kill extracellular bacteria for 24 hours. At the endpoint, the cells were washed twice with DPBS, the cells lysed and scraped in 0.3 mL of lysis solution (40 mg/ml saponin, 8 mL/L polypropylene glycol P-2000, 9.6 μg/ml sodium polyanethol sulfonate) to release intracellular bacteria. The cell lysate was transferred into sterile tubes and 10-fold serially diluted. The number of recovered bacteria was quantified by plating the serial dilutions on BHI agar with rifampicin (100 μg/mL) plates.

### Skin abscess infection mouse model

*E. faecalis* strains were grown in BHI medium to an OD_600nm_ = 0.5 at 37 °C. Next, cells were washed twice with sterile PBS (pH 7.4, 12,000 x g for 2 min) and resuspended to a final concentration of 5×10^7^ colony-forming units (CFU) per mL pre-determined through direct colony counting by CFU plating for each of the strains. Six-week-old female CD-1 mice were anesthetized with isoflurane and had their backs sterilized and shaved. A superficial linear skin abrasion was made with a needle to damage the stratum corneum and upper layer of the epidermis. An aliquot of 20 μL containing the bacterial load resuspended in PBS was inoculated over the scratched area. Animals were euthanized and the area of scarified skin was excised every two days for 10 days, homogenized using a bead beater for 20 minutes (25 Hz), and 10-fold serially diluted for CFU quantification in BHI agar plates with rifampicin (100 μg/mL). The experiments were performed with 4 mice per group^72,73^. All experiments were performed blindly, and no animal subjects were excluded from the analysis. The skin abscess infection mouse model was approved by the University Laboratory Animal Resources (ULAR) from the University of Pennsylvania (Protocol 806763).

## Acknowledgements

We thank members of the Bhatt lab for their helpful discussions and expertise. We thank Matt Gill and Meena Chakraborty for technical assistance. We are grateful to Dr. Christopher Kristich from the Medical College of Wisconsin for sharing cloning reagents to create the deletion mutants in this study, Dr. Michael Federle and Dr. Jennifer Chang for sharing cloning reagents and protocols for experiments with the reporter strains. We thank Dr. Kimberly Kline and Choo Pei Yi for sharing *pur* gene deletion mutants. We thank Dean Felsher for access to the QuantStudio 12K Flex qPCR machine, David Solow-Cordero for assisting in setting up and providing access to the Biomek FX at the Stanford Functional Genomics Facility and High-Throughput Bioscience Center. Biorender was used for creating schematic illustrations in this manuscript. This work was supported by an AACR Fellowship (S.Z.). C.F.N. holds a Presidential Professorship at the University of Pennsylvania and acknowledges funding from the Procter & Gamble Company, United Therapeutics, a BBRF Young Investigator Grant, the Nemirovsky Prize, Penn Health-Tech Accelerator Award, Defense Threat Reduction Agency grants HDTRA11810041 and HDTRA1-23-1-0001, and the Dean’s Innovation Fund from the Perelman School of Medicine at the University of Pennsylvania. Research reported in this publication was supported by the Langer Prize (AIChE Foundation), the NIH R35GM138201, and DTRA HDTRA1-21-1-0014. The LC/MS work was supported by the Mass Spectrometry Core of the Salk Institute with funding from NIH-NCI CCSG: P30 014195 and the Helmsley Center for Genomic Medicine. This work was supported by a National Institutes of Health R01 AI148623 (A.S.B.), National Institutes of Health R01 AI143757 (A.S.B.), Paul Allen Foundation Distinguished Investigator Award (A.S.B.), Stand Up to Cancer Convergence Award (A.S.B.).

## Author contributions

S.Z., H.S., A.S.B. conceived of the study. S.Z., H.S., A.S.B., M.D.T.T., C.F.N. designed experiments, and analyzed data. S.M.B. created deletion mutant and reporter strains for this study and contributed to the biofilm assays. H.S. identified the area of focus for the initial computational search to focus on discovering quorum sensing systems, designed the computational pipeline and conducted the bioinformatics analysis for identifying peptide-based communication systems in HMP reference genomes. M.D.T.T. designed, conducted, and analyzed the murine infection model experiments. C.F.N. advised on the design of the murine infection model experiments. J.V. performed the peptide enrichment and prepared the samples for targeted LC/MS analysis. A.F.M.P. performed LC/MS analysis for peptide detection and quantification and analyzed the data. D.F. advised on the expression and purification of recombinant peptide receptors in complex to smORFs. A.S. advised on the design of the peptide enrichment and LC/MS analysis experiments. Y.P. generated the phylogenetic tree. H.R. designed experiments. S.Z. and A.S.B prepared the first draft, reviewed and edited the manuscript. All authors read and approved the final manuscript and take responsibility for its content.

## SUPPLEMENTARY INFORMATION

### 1. Supplementary Tables

Supplementary Table 1. Putative RRNPP systems identified in HMP reference genomes

Supplementary Table 2. Putative RRNPP systems in enterococcal species

Supplementary Table 3. Differentially expressed genes in response to treatment with NAPHQ

Supplementary Table 4. Differentially expressed genes in response to treatment with NVPHV

Supplementary Table 5. Primers used in this study

### 2. Supplementary Figures

### 3. Supplementary Data

Sequencing data

**Supplementary Figure 1.**
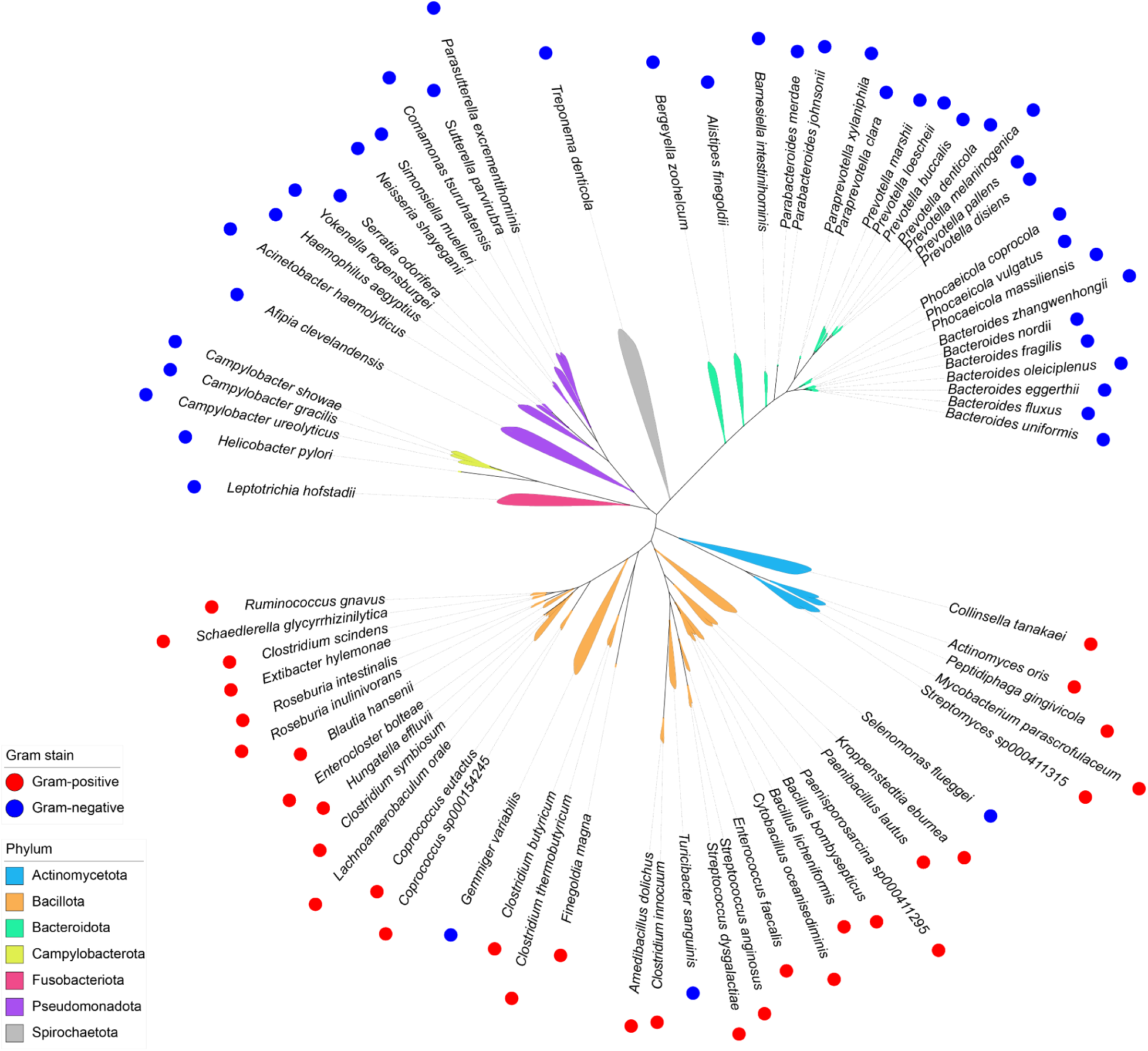
Phylogenetic tree of the HMP bacterial species that encode putative RRNPP systems. Phylogenetic tree of HMP reference genomes representing the 75 bacterial species in which putative RRNPP systems were identified. The tree was built using GTDB-Tk^1^. The species are colored based on their phylum, and labeled based on their Gram stain status.

**Supplementary Figure 2.**
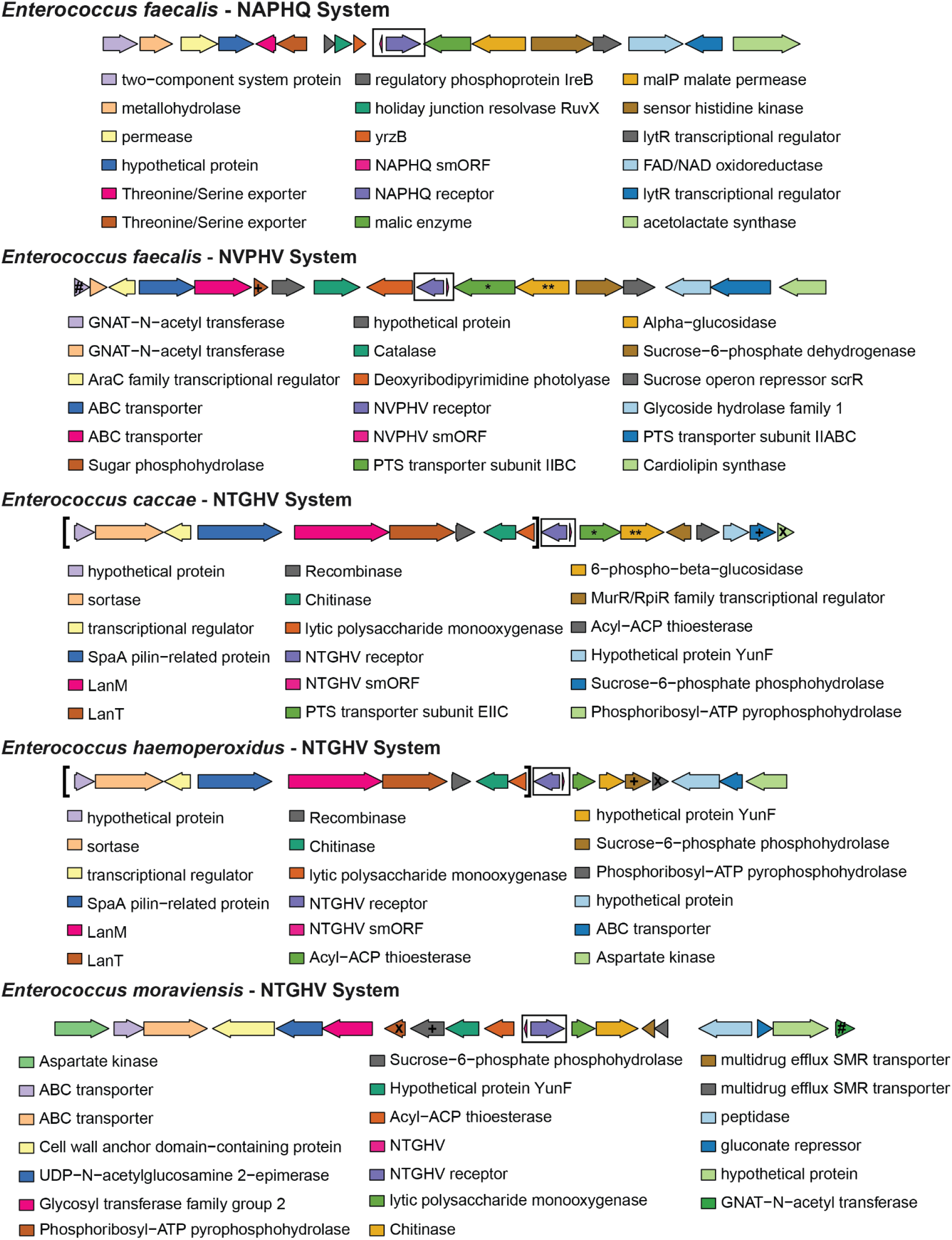
Analysis of the genomic neighborhoods of the putative RRNPP systems in *Enterococcus spp*. The genomes of the enterococcal species that have putative RRNPP systems similar to the NAPHQ and NVPHV systems (*E. caccae, E. haemoperoxidus*, and *E. moraviensis*) were functionally annotated using eggNOG-mapper^2^. The genomic regions comprising 7-9 genes upstream and downstream of the putative RRNPP systems are illustrated with the RRNPP systems (smORF + receptor) shown in a black rectangle. When compared to the genomic regions around the NAPHQ and NVPHV systems in *E. faecalis*, the regions around the NTGHV system in *E. caccae, E. haemoperoxidus*, and *E. moraviensis* share a number of conserved features with the NVPHV system in *E. faecalis*, namely sugar transporter or sugar utilization genes. Specifically, similar to the NVPHV system in *E. faecalis*, *E. caccae* has a phosphotransferase system (PTS) transporter subunit (labeled *) and a sugar glucosidase (labeled **) next to its NTGHV system. A sugar phosphohydrolase (labeled +) is present in the genomic context of the NVPHV system in *E. faecalis* and the NTGHV systems of the three other enterococcal species. Finally, both *E. faecalis* and *E. moraviensis* have a GNAT-N-acetyl transferase (labeled #) in the genomic neighborhood of their NVPVH and NTGHV systems, respectively. In contrast, the NTGHV systems in all three species don’t share any features in common with the NAPHQsystem in *E. faecalis*.

**Supplementary Figure 3.**
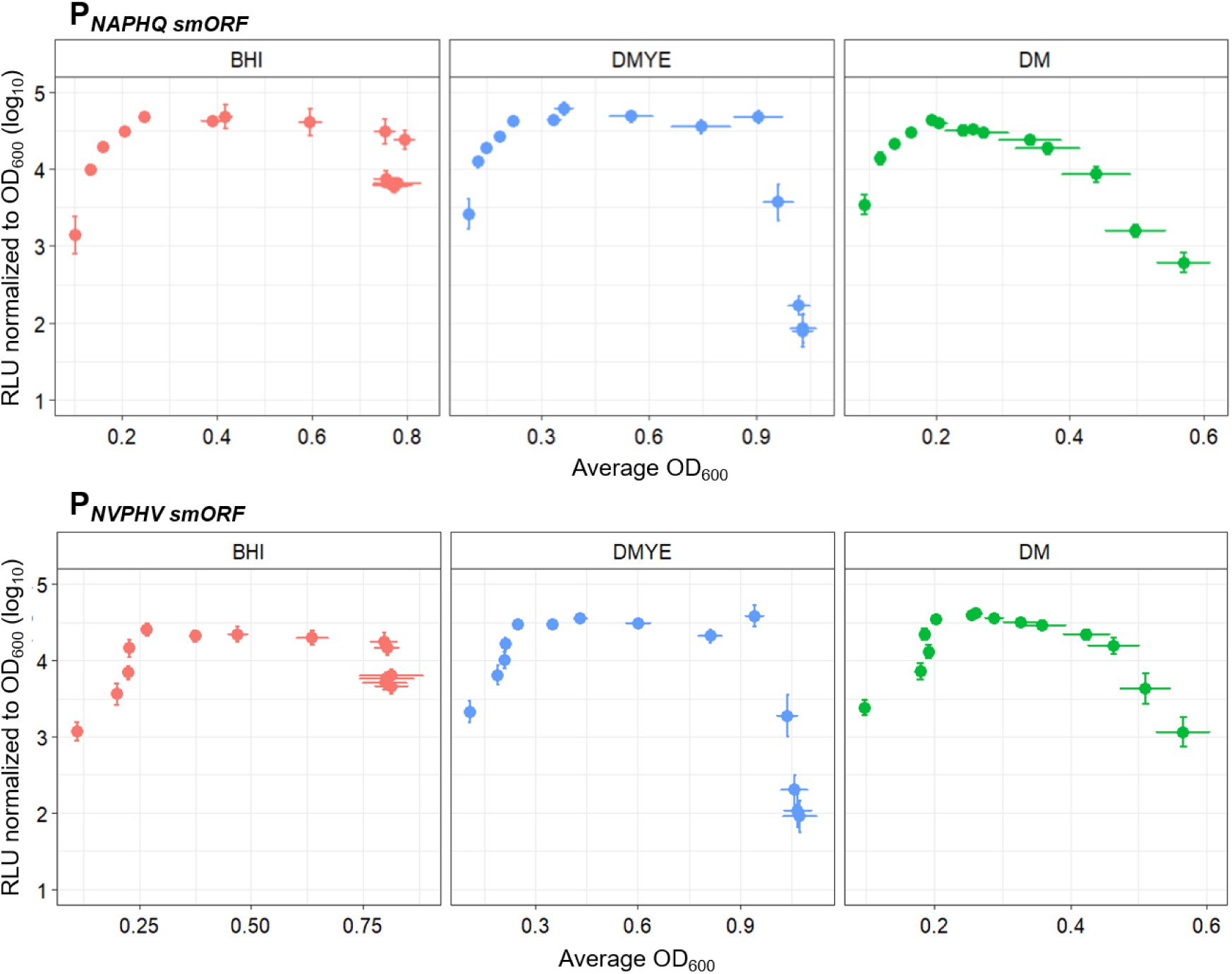
smORF gene promoter activity is cell density-dependent. Transcriptional activity of the *smORF* gene promoters was measured in different media using a promoter-luciferase reporter plasmid system in *E. faecalis*. *E. faecalis* strains (n = 4 biological replicates) harboring P*_NAPHQ smORF_*-*luxABCDE* or P*_NVPHV smORF_*-*luxABCDE* reporter vectors were grown in in BHI, defined medium (DM), or defined medium supplemented with 0.2% yeast extract (DMYE). Luciferase expression and optical density (OD_600_) were measured over time. Data are expressed as the mean of n = 4 biological replicates ± standard deviation.

**Supplementary Figure 4.**
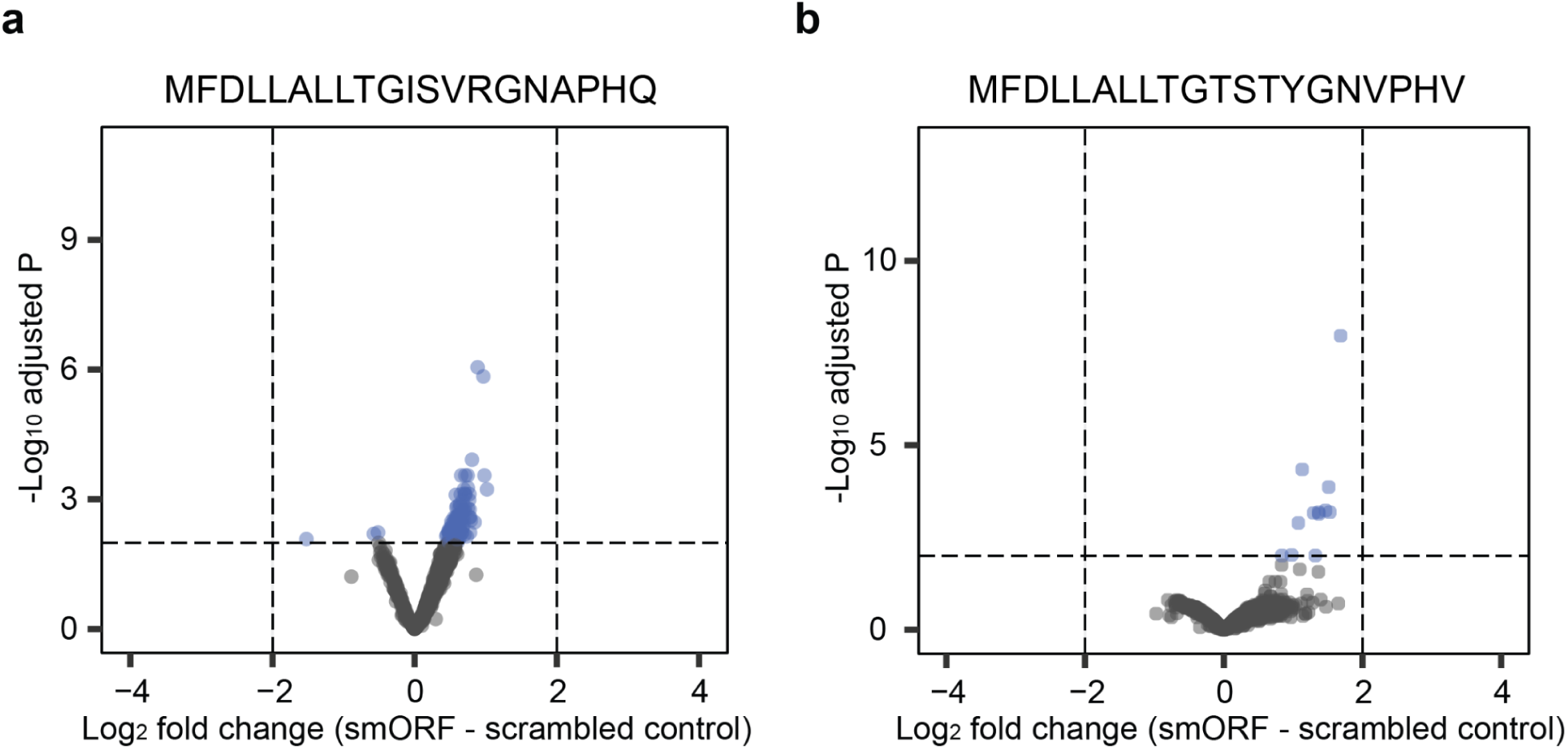
The full-length signaling smORFs do not elicit a significant transcriptional response in *E. faecalis*. Volcano plots gene expression analysis of WT *E. faecalis* grown in DM and treated with 5 μM of the full-length 20 amino acid smORFs, **(a)** NAPHQ full-length smORF and **(b)** NVPHV full-length smORF, or their respective scrambled controls for 15 min before the samples were collected, quenched, and the RNA extracted for analysis. Differentially expressed genes are defined as those that display at least 4-fold change in gene expression relative to the scrambled control with FDR cutoff < 0.01.

**Supplementary Figure 5.**
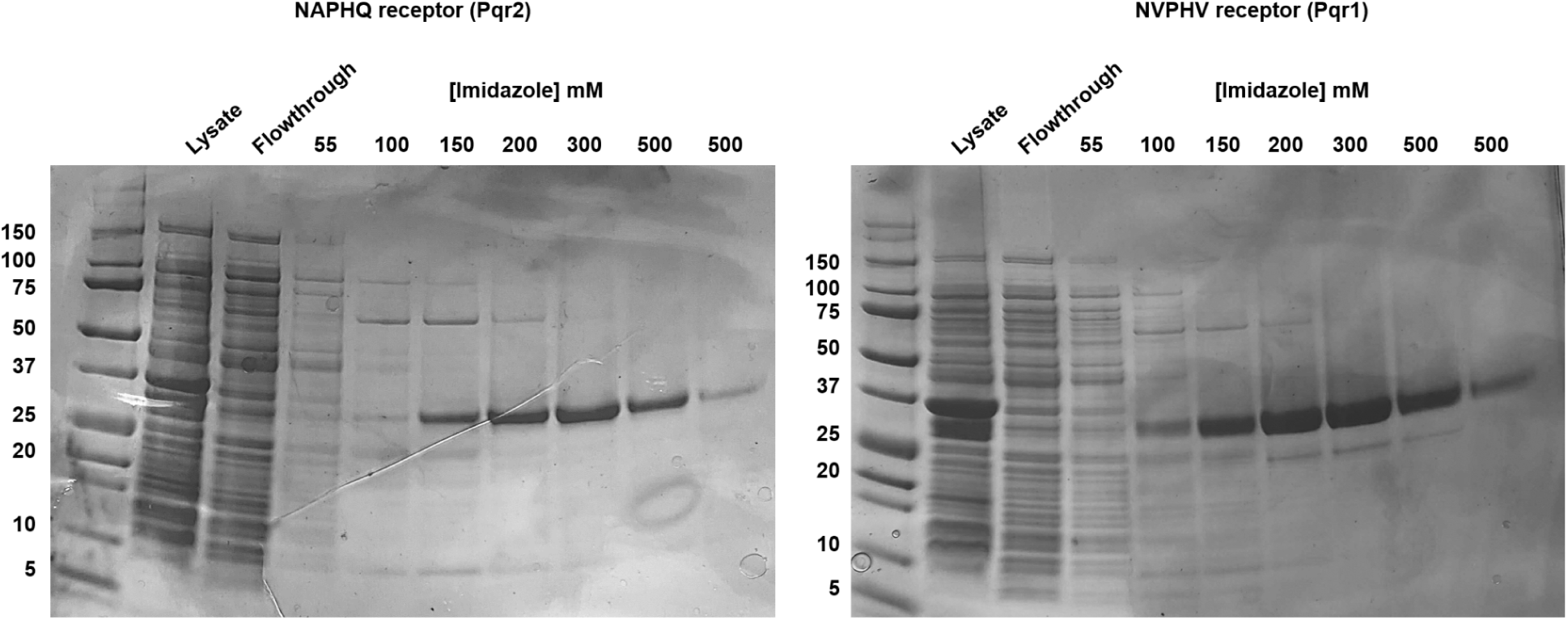
SDS-PAGE analysis of recombinant smORF receptors purification. Recombinant 6xHis-tag labeled smORF receptor proteins were purified by nickel affinity chromatography. Purification was done by gravity flow by placing clarified lysate nickel resin mix in the column and allowing it to flow through. Bound protein was washed with increasing concentrations of imidazole. Fractions were analyzed on SDS-PAGE tricine gels. The purest fractions (eluted with 500 mM imidazole) were pooled, buffer exchanged, and used in experiments.

## REFERENCES

1. Hardman, A. M., Stewart, G. S. & Williams, P. Quorum sensing and the cell-cell communication dependent regulation of gene expression in pathogenic and non-pathogenic bacteria. Antonie Van Leeuwenhoek 74, 199–210 (1998).

2. Waters, C. M. & Bassler, B. L. Quorum sensing: cell-to-cell communication in bacteria. Annu. Rev. Cell Dev. Biol. 21, 319–346 (2005).

3. Declerck, N. et al. Structure of PlcR: Insights into virulence regulation and evolution of quorum sensing in Gram-positive bacteria. Proc. Natl. Acad. Sci. U. S. A. 104, 18490– 18495 (2007).

4. Winson, M. K. et al. Multiple N-acyl-L-homoserine lactone signal molecules regulate production of virulence determinants and secondary metabolites in Pseudomonas aeruginosa. Proc. Natl. Acad. Sci. U. S. A. 92, 9427–9431 (1995).

5. Parsek, M. R. & Greenberg, E. P. Acyl-homoserine lactone quorum sensing in gram-negative bacteria: a signaling mechanism involved in associations with higher organisms. Proc. Natl. Acad. Sci. U. S. A. 97, 8789–8793 (2000).

6. Flemming, H.-C. et al. Biofilms: an emergent form of bacterial life. Nat. Rev. Microbiol. 14, 563–575 (2016).

7. Daniels, R., Vanderleyden, J. & Michiels, J. Quorum sensing and swarming migration in bacteria. FEMS Microbiol. Rev. 28, 261–289 (2004).

8. Dubois, T. et al. Activity of the Bacillus thuringiensis NprR-NprX cell-cell communication system is co-ordinated to the physiological stage through a complex transcriptional regulation. Mol. Microbiol. 88, 48–63 (2013).

9. Jiang, M., Grau, R. & Perego, M. Differential processing of propeptide inhibitors of Rap phosphatases in Bacillus subtilis. J. Bacteriol. 182, 303–310 (2000).

10. Hirt, H. et al. Enterococcus faecalis Sex Pheromone cCF10 Enhances Conjugative Plasmid Transfer In Vivo. MBio 9, (2018).

11. Dunny, G. M. & Berntsson, R. P.-A. Enterococcal Sex Pheromones: Evolutionary Pathways to Complex, Two-Signal Systems. J. Bacteriol. 198, 1556–1562 (2016).

12. Shanker, E. & Federle, M. J. Quorum Sensing Regulation of Competence and Bacteriocins in Streptococcus pneumoniae and mutans. Genes 8, (2017).

13. Mok, K. C., Wingreen, N. S. & Bassler, B. L. Vibrio harveyi quorum sensing: a coincidence detector for two autoinducers controls gene expression. EMBO J. 22, 870–881 (2003).

14. Maldonado-Barragán, A. & West, S. A. The cost and benefit of quorum sensing-controlled bacteriocin production in Lactobacillus plantarum. J. Evol. Biol. 33, 101–111 (2020).

15. Meng, F. et al. Acetate Activates Lactobacillus Bacteriocin Synthesis by Controlling Quorum Sensing. Appl. Environ. Microbiol. 87, e0072021 (2021).

16. Duval, M. & Cossart, P. Small bacterial and phagic proteins: an updated view on a rapidly moving field. Curr. Opin. Microbiol. 39, 81–88 (2017).

17. Storz, G., Wolf, Y. I. & Ramamurthi, K. S. Small proteins can no longer be ignored. Annu. Rev. Biochem. 83, 753–777 (2014).

18. Plaza, S., Menschaert, G. & Payre, F. In Search of Lost Small Peptides. Annu. Rev. Cell Dev. Biol. 33, 391–416 (2017).

19. Su, M., Ling, Y., Yu, J., Wu, J. & Xiao, J. Small proteins: untapped area of potential biological importance. Front. Genet. 4, 286 (2013).

20. Sberro, H. et al. Large-Scale Analyses of Human Microbiomes Reveal Thousands of Small, Novel Genes. Cell 178, 1245–1259.e14 (2019).

21. Fesenko, I., Sahakyan, H., Shabalina, S. A. & Koonin, E. V. The Cryptic Bacterial Microproteome. *bioRxiv* 2024.02.17.580829 (2024) doi:10.1101/2024.02.17.580829.

22. Bartholomäus, A. et al. smORFer: a modular algorithm to detect small ORFs in prokaryotes. Nucleic Acids Res. 49, e89 (2021).

23. Durrant, M. G. & Bhatt, A. S. Automated Prediction and Annotation of Small Open Reading Frames in Microbial Genomes. Cell Host Microbe 29, 121–131.e4 (2021).

24. Monnet, V., Juillard, V. & Gardan, R. Peptide conversations in Gram-positive bacteria. Crit. Rev. Microbiol. 42, 339–351 (2016).

25. Neiditch, M. B., Capodagli, G. C., Prehna, G. & Federle, M. J. Genetic and Structural Analyses of RRNPP Intercellular Peptide Signaling of Gram-Positive Bacteria. Annu. Rev. Genet. 51, 311–333 (2017).

26. Perez-Pascual, D., Monnet, V. & Gardan, R. Bacterial Cell-Cell Communication in the Host via RRNPP Peptide-Binding Regulators. Front. Microbiol. 7, 706 (2016).

27. Parashar, V., Mirouze, N., Dubnau, D. A. & Neiditch, M. B. Structural basis of response regulator dephosphorylation by Rap phosphatases. PLoS Biol. 9, e1000589 (2011).

28. Bongiorni, C., Ishikawa, S., Stephenson, S., Ogasawara, N. & Perego, M. Synergistic regulation of competence development in Bacillus subtilis by two Rap-Phr systems. J. Bacteriol. 187, 4353–4361 (2005).

29. Gohar, M. et al. The PlcR virulence regulon of Bacillus cereus. PLoS One 3, e2793 (2008).

30. Dubois, T. et al. Necrotrophism is a quorum-sensing-regulated lifestyle in Bacillus thuringiensis. PLoS Pathog. 8, e1002629 (2012).

31. Rocha, J. et al. Evolution and some functions of the NprR-NprRB quorum-sensing system in the Bacillus cereus group. Appl. Microbiol. Biotechnol. 94, 1069–1078 (2012).

32. Underhill, S. A. M. et al. Intracellular Signaling by the comRS System in Streptococcus mutans Genetic Competence. mSphere 3, (2018).

33. Lebreton, F. et al. Tracing the Enterococci from Paleozoic Origins to the Hospital. Cell 169, 849–861.e13 (2017).

34. Singh, R. P. & Nakayama, J. Quorum-Sensing Systems in Enterococci. Quorum Sensing vs Quorum Quenching: A Battle with No End in Sight 155–163 Preprint at 10.1007/978-81-322-1982-8_14 (2015).

35. Chang, J. C., LaSarre, B., Jimenez, J. C., Aggarwal, C. & Federle, M. J. Two group A streptococcal peptide pheromones act through opposing Rgg regulators to control biofilm development. PLoS Pathog. 7, e1002190 (2011).

36. Rahbari, K. M., Chang, J. C. & Federle, M. J. A Streptococcus Quorum Sensing System Enables Suppression of Innate Immunity. MBio 12, (2021).

37. Wilkening, R. V., Langouët-Astrié, C., Severn, M. M., Federle, M. J. & Horswill, A. R. Identifying genetic determinants of Streptococcus pyogenes-host interactions in a murine intact skin infection model. Cell Rep. 42, 113332 (2023).

38. Zhang, J. Evolution by gene duplication: an update. Trends Ecol. Evol. 18, 292–298 (2003).

39. Conant, G. C. & Wolfe, K. H. Turning a hobby into a job: how duplicated genes find new functions. Nat. Rev. Genet. 9, 938–950 (2008).

40. Andersson, D. I. & Hughes, D. Gene amplification and adaptive evolution in bacteria. Annu. Rev. Genet. 43, 167–195 (2009).

41. Serres, M. H., Kerr, A. R. W., McCormack, T. J. & Riley, M. Evolution by leaps: gene duplication in bacteria. Biol. Direct 4, 46 (2009).

42. Human Microbiome Jumpstart Reference Strains Consortium et al. A catalog of reference genomes from the human microbiome. Science 328, 994–999 (2010).

43. Gélinas, M., Museau, L., Milot, A. & Beauregard, P. B. The de novo Purine Biosynthesis Pathway Is the Only Commonly Regulated Cellular Pathway during Biofilm Formation in TSB-Based Medium in Staphylococcus aureus and Enterococcus faecalis. Microbiol Spectr 9, e0080421 (2021).

44. Gentry-Weeks, C. R., Karkhoff-Schweizer, R., Pikis, A., Estay, M. & Keith, J. M. Survival of Enterococcus faecalis in mouse peritoneal macrophages. Infect. Immun. 67, 2160–2165 (1999).

45. Goncheva, M. I., Flannagan, R. S. & Heinrichs, D. E. De Novo Purine Biosynthesis Is Required for Intracellular Growth of Staphylococcus aureus and for the Hypervirulence Phenotype of a purR Mutant. Infect. Immun. 88, (2020).

46. Goncheva, M. I., Chin, D. & Heinrichs, D. E. Nucleotide biosynthesis: the base of bacterial pathogenesis. Trends Microbiol. 30, 793–804 (2022).

47. Bernard, C., Li, Y., Lopez, P. & Bapteste, E. Large-Scale Identification of Known and Novel RRNPP Quorum-Sensing Systems by RRNPP_Detector Captures Novel Features of Bacterial, Plasmidic, and Viral Coevolution. Mol. Biol. Evol. 40, (2023).

48. Tan, C. A. Z. et al. Purine and carbohydrate availability drive Enterococcus faecalis fitness during wound and urinary tract infections. MBio 15, e0238423 (2024).

49. McDonough, K. A. & Rodriguez, A. The myriad roles of cyclic AMP in microbial pathogens: from signal to sword. Nat. Rev. Microbiol. 10, 27–38 (2011).

50. Irving, S. E., Choudhury, N. R. & Corrigan, R. M. The stringent response and physiological roles of (pp)pGpp in bacteria. Nat. Rev. Microbiol. 19, 256–271 (2021).

51. Jenal, U., Reinders, A. & Lori, C. Cyclic di-GMP: second messenger extraordinaire. Nat. Rev. Microbiol. 15, 271–284 (2017).

52. Martins, F. H. et al. Enterococcus faecalis-derived adenine enhances enterohaemorrhagic Escherichia coli Type 3 Secretion System-dependent virulence. Nat Microbiol (2024) doi:10.1038/s41564-024-01747-1.

53. Monteagudo-Cascales, E. et al. Ubiquitous purine sensor modulates diverse signal transduction pathways in bacteria. Nat. Commun. 15, 5867 (2024).

54. Käll, L., Krogh, A. & Sonnhammer, E. L. L. A combined transmembrane topology and signal peptide prediction method. J. Mol. Biol. 338, 1027–1036 (2004).

55. Karpenahalli, M. R., Lupas, A. N. & Söding, J. TPRpred: a tool for prediction of TPR-, PPR- and SEL1-like repeats from protein sequences. BMC Bioinformatics 8, 2 (2007).

56. Fu, L., Niu, B., Zhu, Z., Wu, S. & Li, W. CD-HIT: accelerated for clustering the next-generation sequencing data. Bioinformatics 28, 3150–3152 (2012).

57. Chaumeil, P.-A., Mussig, A. J., Hugenholtz, P. & Parks, D. H. GTDB-Tk: a toolkit to classify genomes with the Genome Taxonomy Database. Bioinformatics 36, 1925–1927 (2019).

58. Letunic, I. & Bork, P. Interactive Tree of Life (iTOL) v6: recent updates to the phylogenetic tree display and annotation tool. Nucleic Acids Res. 52, W78–W82 (2024).

59. Kristich, C. J., Chandler, J. R. & Dunny, G. M. Development of a host-genotype-independent counterselectable marker and a high-frequency conjugative delivery system and their use in genetic analysis of Enterococcus faecalis. Plasmid 57, 131–144 (2007).

60. Vesić, D. & Kristich, C. J. A Rex family transcriptional repressor influences H2O2 accumulation by Enterococcus faecalis. J. Bacteriol. 195, 1815–1824 (2013).

61. Kellogg, S. L., Little, J. L., Hoff, J. S. & Kristich, C. J. Requirement of the CroRS Two-Component System for Resistance to Cell Wall-Targeting Antimicrobials in Enterococcus faecium. Antimicrob. Agents Chemother. 61, (2017).

62. Gibson, D. G. et al. Enzymatic assembly of DNA molecules up to several hundred kilobases. Nat. Methods 6, 343–345 (2009).

63. Cruz-Rodz, A. L. & Gilmore, M. S. High efficiency introduction of plasmid DNA into glycine treated Enterococcus faecalis by electroporation. Mol. Gen. Genet. 224, 152–154 (1990).

64. Dunny, G. M., Lee, L. N. & LeBlanc, D. J. Improved electroporation and cloning vector system for gram-positive bacteria. Appl. Environ. Microbiol. 57, 1194–1201 (1991).

65. MacLean, B., et al. Skyline: an open source document editor for creating and analyzing targeted proteomics experiments. Bioinformatics 26, 966–968 (2010).

66. LaSarre, B., Chang, J. C. & Federle, M. J. Redundant group a streptococcus signaling peptides exhibit unique activation potentials. J. Bacteriol. 195, 4310–4318 (2013).

67. Chang, J. C. & Federle, M. J. PptAB Exports Rgg Quorum-Sensing Peptides in Streptococcus. PLoS One 11, e0168461 (2016).

68. Krueger, F. TrimGalore: A Wrapper around Cutadapt and FastQC to Consistently Apply Adapter and Quality Trimming to FastQ Files, with Extra Functionality for RRBS Data. (Github).

69. Applied Research Press. Ultrafast and Memory-Efficient Alignment of Short DNA Sequences to the Human Genome. (CreateSpace Independent Publishing Platform, 2015).

70. Quinlan, A. R. & Hall, I. M. BEDTools: a flexible suite of utilities for comparing genomic features. Bioinformatics 26, 841–842 (2010).

71. Love, M. I., Huber, W. & Anders, S. Moderated estimation of fold change and dispersion for RNA-seq data with DESeq2. Genome Biol. 15, 550 (2014).

72. Torres, M. D. T. et al. Structure-function-guided exploration of the antimicrobial peptide polybia-CP identifies activity determinants and generates synthetic therapeutic candidates. Commun Biol 1, 221 (2018).

73. Silva, O. N. et al. Repurposing a peptide toxin from wasp venom into antiinfectives with dual antimicrobial and immunomodulatory properties. Proc. Natl. Acad. Sci. U. S. A. 117, 26936–26945 (2020).

## REFERENCES

1. Chaumeil, P.-A., Mussig, A. J., Hugenholtz, P. & Parks, D. H. GTDB-Tk: a toolkit to classify genomes with the Genome Taxonomy Database. Bioinformatics 36, 1925–1927 (2019).

2. Cantalapiedra, C. P., Hernández-Plaza, A., Letunic, I., Bork, P. & Huerta-Cepas, J. eggNOG-mapper v2: Functional Annotation, Orthology Assignments, and Domain Prediction at the Metagenomic Scale. Mol. Biol. Evol. 38, 5825–5829 (2021).

